# A giant virus awakens polinton-like virophages in the green alga *Tetraselmis*, revealing an inducible antiviral defense system

**DOI:** 10.1101/2025.10.09.676808

**Authors:** Sonia Bouchard, Malik Blanc, Jean-Pierre Baudoin, Julien Andreani, Bernard La Scola, Guillaume Blanc

## Abstract

Giant double-stranded DNA viruses profoundly influence algal population dynamics, yet their interactions with co-infecting mobile elements remain poorly understood. Here, we describe a natural tripartite infection system in the marine unicellular alga *Tetraselmis striata*, where a newly isolated giant virus, *Oceanusvirus lionense* (TetV-2), triggers the productive replication of multiple polinton-like viruses (PLVs). One PLV (Tsv-S2b) was co-isolated from the same seawater sample, while two others (Tsv-S2a and Tsv-S3b) derive from endogenous elements integrated in the algal genome. These PLVs depend strictly on TetV-2 for propagation and exert a virophage-like effect, reducing TetV-2 yields in a dose-dependent manner. Comparative genomics shows that virophagic PLVs retain a conserved structural module but harbor divergent replication and integration genes, supporting modular evolution of the PLV lineage. The reactivation of integrated PLVs mirrors the behavior of endogenous virophages in *Cafeteria burkhardae*, suggesting that specific Oceanusvirus–PLV compatibilities govern reactivation. This work provides direct evidence that integrated PLVs in green algae can transition to a virophage lifestyle upon infection by a giant virus, highlighting an inducible antiviral defense mechanism within photosynthetic protists.

**Author Summary:** Marine microalgae are frequently infected by giant viruses, but how these infections are regulated is poorly understood. In the green alga *Tetraselmis striata*, we discovered that a giant virus can “wake up” small DNA viruses known as polinton-like viruses (PLVs) that are normally integrated and silent within the algal genome. Once activated, these PLVs multiply using the giant virus’s replication machinery and, in turn, reduce its production. This helper-dependent relationship mirrors the behavior of virophages previously known only in non-photosynthetic protists. Our results reveal that microalgae possess an inducible antiviral system based on the reactivation of endogenous viral elements, suggesting that such interactions may be widespread in the oceans and play an important role in controlling algal virus epidemics.

## Introduction

Giant double-stranded DNA (dsDNA) viruses of the phylum Nucleocytoviricota are major regulators of microbial life in aquatic ecosystems [1–3]. By infecting and lysing abundant microalgae and protists, these viruses influence primary production, nutrient cycling, and bloom dynamics across marine environments [4–6]. Beyond their ecological importance, giant viruses are central to the evolution of eukaryotic cells, acting as reservoirs and vectors of metabolic and informational genes [7,8]. Yet, these viral pathogens may not act alone: many interact with smaller mobile genetic elements, such as virophages and polinton-like viruses (PLVs), that can alter the outcome of infection in profound ways.

Virophages (class ‘Virophaviricetes’) are small dsDNA viruses that replicate only in the presence of a co-infecting giant virus [9–12]. They parasitize the giant virus’s replication factory, diverting resources to produce their own progeny while reducing the yield and infectivity of the helper virus. The prototypical virophage Mavirus, discovered in the heterotrophic flagellate *Cafeteria roenbergensis*, provided the first demonstration that such elements can also integrate into the host nuclear genome [13,14]. Upon subsequent infection by the giant virus CroV, integrated Mavirus elements are transcriptionally reactivated and produce virions that inhibit CroV replication [13,15]. This reactivation depends on the CroV late transcriptional machinery and is abolished by inhibitors of viral or host protein synthesis, establishing a mechanistic link between helper-virus gene expression and provirophage induction. The Cafeteria–CroV–Mavirus model thus revealed a striking example of an inducible, virus-triggered antiviral defense system in a unicellular eukaryote.

Polinton-like viruses (class ‘Aquintoviricetes’) occupy a related but broader evolutionary niche at the interface between transposable elements and bona fide viruses [11,12,16]. They share with virophages a conserved structural module, including major and minor capsid proteins of the double jelly-roll type and a DNA-packaging ATPase. PLVs have been identified as integrated elements within the nuclear genomes of a wide range of eukaryotic lineages, from unicellular protists to multicellular animals, and are particularly abundant in aquatic environments, where they also occur as free viral particles [17–21]. In some microalgae, PLVs occur both as autonomous lytic viruses and as integrated elements, implying dual lifestyles that may alternate between vertical inheritance and horizontal propagation [22,23]. Despite their ubiquity, however, no direct experimental evidence had previously demonstrated PLV reactivation by a giant virus or their capacity to behave as virophages.

The green algal genus Tetraselmis (Chlorodendrophyceae) provides an ideal system to explore these relationships. Tetraselmis species are globally distributed coastal microalgae that contribute significantly to marine productivity and occasionally form dense blooms [24–26]. They are natural hosts to both giant viruses and PLVs. The first characterized giant virus of this group, Oceanusvirus kaneohense (TetV-1), infects an unidentified Tetraselmis strain and encodes a large circular genome exceeding 650 kb [27]. Separately, the small dsDNA virus Tetraselmis striata virus N1 (TsV-N1) defines a clade of PLVs capable of autonomous lytic infection [22,28]. Comparative genomics has shown that multiple Tetraselmis genomes harbor dozens of integrated viral elements related to both TetV-like giant viruses and TsV-like PLVs, suggesting recurrent endogenization and potential interactions between these lineages [23]. However, whether these integrated PLVs can be reactivated and whether such events influence giant-virus infection dynamics have remained open questions.

Here, we report the discovery of a natural tripartite infection system in *T. striata*, in which a newly isolated giant virus, *Oceanusvirus lionense* (TetV-2), triggers the productive replication of multiple PLVs. One PLV, Tsv-S2b, was co-isolated from the same seawater sample, while at least two additional PLVs (Tsv-S2a and Tsv-S3b) correspond to elements integrated within the algal genome. Using a combination of time-resolved infection assays, electron microscopy, and quantitative PCR, we demonstrate that these PLVs replicate exclusively in the presence of TetV-2, emerge after the onset of giant virus replication, and suppress TetV-2 yields in a dose-dependent manner, key features of a virophage-like lifestyle. Comparative genomic analysis reveals that all *Tetraselmis* PLVs share a conserved structural module but differ in replication and integration genes, supporting a model of modular evolution that enables transitions between autonomous, integrated, and helper-dependent lifestyles. This system provides the first direct evidence that integrated PLVs in *Tetraselmis* can be awakened by a giant virus, bridging the gap between endogenous viral elements and virophages. Beyond expanding the known repertoire of virophage-like interactions in marine microalgae, our findings have broader implications for understanding the evolutionary and ecological consequences of virus–virus interactions, the genomic plasticity of eukaryotic microorganisms in marine ecosystems and the ancient origins of antiviral defense systems in unicellular life.

## Results

### Isolation and genome of a new Tetraselmis giant virus

A culture of *Tetraselmis striata* was inoculated with 0.45 µm–filtered environmental water collected from the Lion Lagoon (southern France), resulting in complete culture lysis. During the infection process, cells exhibited a marked loss of motility and underwent deflagellation, accompanied by a pronounced decline in viable cell numbers as observed microscopically. To assess the reproducibility and stability of the lytic activity, and to fulfill Koch’s postulates, several successive infection cycles were performed by inoculating fresh *T. striata* cultures with 0.45 µm–filtered lysates derived from the preceding lysed cultures.

Diagnostic short-read sequencing of the initial 0.45 µm–filtered lysate yielded 20 contigs encoding proteins predominantly most similar to homologs from *Oceanusvirus kaneohense* (TetV-1). These contigs spanned a total of 632 kb and displayed a relatively uniform sequencing depth of 792× ± 90. The proteins encoded by these contigs matched distinct, non-overlapping regions of the TetV-1 genome, indicating that they likely represent fragments of a single virus genome closely related to TetV-1. However, attempts to reconstruct a complete genome from the short-read dataset, including multiple contig extension trials, were unsuccessful.

To overcome this limitation, we isolated a single viral clone from the lysate, designated TetV-2, through several rounds of serial dilution to extinction, and sequenced its genome using a long-read approach. Assembly produced a single circular contig of 660,680 bp, encoding 640 predicted proteins and 4 tRNAs. A large fraction of these proteins (n = 389; 61%) shared significant similarity with TetV-1 homologs, of which 348 had their best BLAST match in TetV-1 (**Fig. 1A**). Reciprocal best-hit analysis identified 343 orthologous pairs. The four TetV-2 tRNAs corresponded to a subset of the TetV-1 tRNA repertoire (10 genes) and, as in TetV-1, were clustered within a single genomic locus.

**Figure 1:**
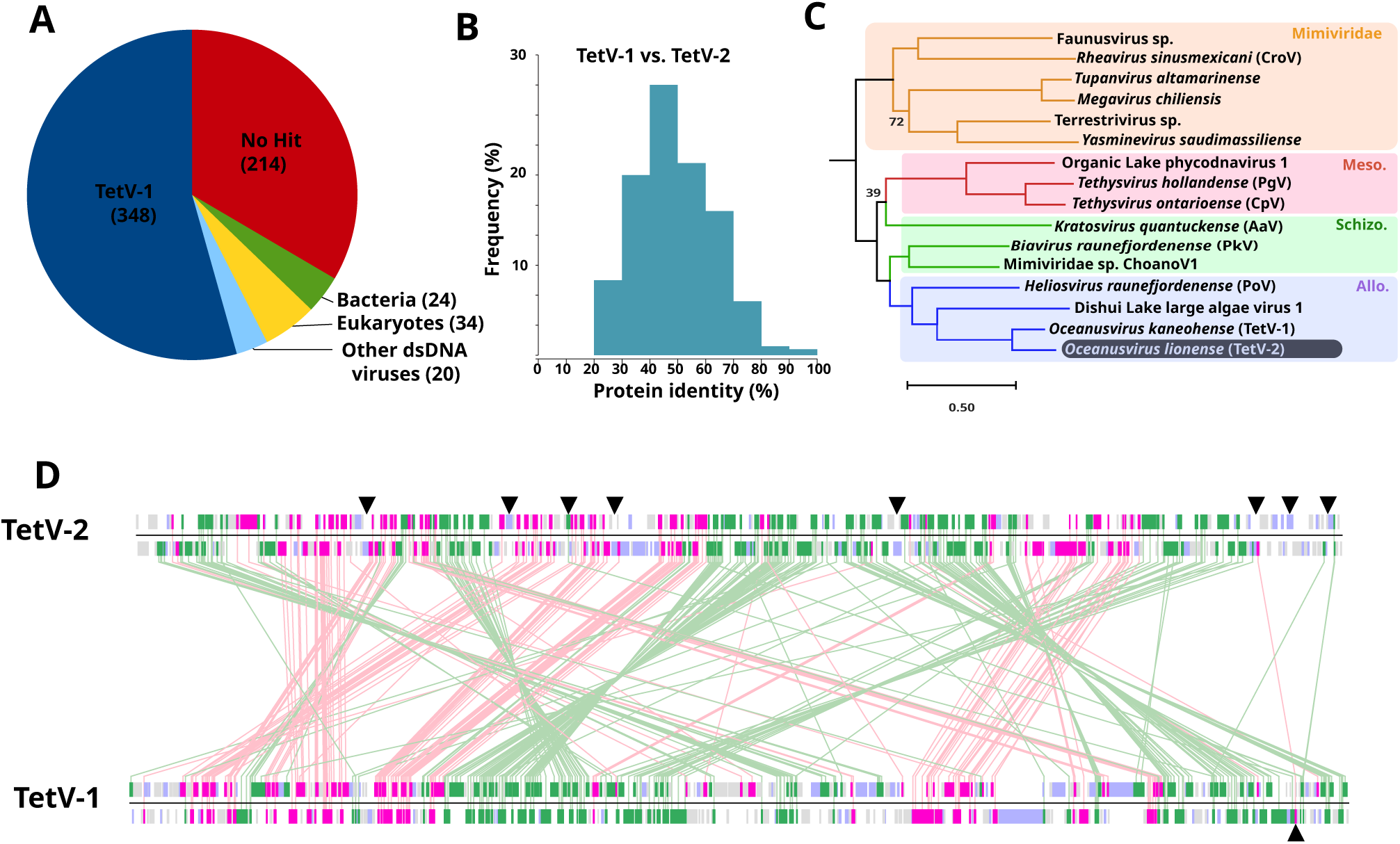
Comparative genomics of TetV. (A) Taxonomic origin of the best matches in the GenBank NR database to TetV-2 proteins.(B) Sequence identity distribution among reciprocal best matches between TetV-1 and TetV-2 proteins.(C) Maximum likelihood phylogenetic reconstruction (IQ-TREE) based on a concatenated alignment of the A18 helicase, packaging ATPase, and DNA polymerase. The tree was rooted using chlorovirus and mamiellovirus homologs as outgroup sequences (not shown). Bootstrap support values <90% are indicated.(D) Gene colinearity between TetV-1 and TetV-2 genomes. Genes are represented as colored boxes: pink, conserved genes on the same strand; green, conserved genes on opposite strands; purple, genes with significant NR matches but lacking reciprocal best hits between the two viruses; gray, hypothetical genes without significant matches. Triangles shows gene doublets encoding a bacterial transposase-related protein and an uncharacterized “transposase-associated” protein.

TetV-2 lacked several genes present in TetV-1, including two fermentation-related genes encoding pyruvate formate-lyase (TetV_428) and its activating enzyme (TetV_456) [27], as well as a photolyase (TetV_298) and a deoxyribodipyrimidine photolyase-related protein (TetV_522), both likely involved in UV-induced DNA repair. Notably, TetV-2 also lacked thymidylate synthase (TetV_240) and thymidylate kinase (TetV_225), which catalyze successive steps in the conversion of dUMP to dTDP. In contrast, 54 TetV-2 proteins had no detectable similarity to TetV-1 but did display significant matches to bacterial, eukaryotic, or viral proteins. Among these were enzymes involved in lipid metabolism, including a very long-chain fatty acid elongase (TetV2_00470) and a lysophospholipid acyltransferase (TetV2_00549). We also identified two genes encoding putative mitochondrial proteins: an “altered inheritance of mitochondria” protein 24 (AIM24; TetV2_00382) and a calcium-binding mitochondrial carrier protein (TetV2_00259). A substantial fraction of TetV-2 proteins (n = 264; 41%) had no detectable homologs in the NR database; these were globally shorter than conserved proteins (average length 200 vs. 377 amino acids, respectively). On average, orthologous proteins shared 49% sequence identity at the amino acid level (**Fig. 1B**).

Phylogenetic placement of TetV-2 within the Allomimiviridae [29] was supported by analyses based on concatenated alignments of three hallmark Megaviricetes proteins: the A18 helicase, packaging ATPase, and DNA polymerase (**Fig. 1C**). Despite infecting distinct *Tetraselmis* hosts, TetV-1 and TetV-2 were more closely related to one another than to any endogenous TetV-like elements integrated within *Tetraselmis* genomes, including those of *T. striata*, the host of TetV-2 (**Fig. S1**). This indicates that the diversity of *Tetraselmis*-infecting viruses extends beyond that represented by TetV-1 and TetV-2.

Base composition differed substantially between the two viruses: TetV-1 and TetV-2 had GC contents of 41.2% and 57.8%, respectively. Even higher GC levels were observed among endogenous TetV-like elements in *T. striata*, *T. suecica*, and *T. chui* (ranging from 58.8% to 73.9%). These compositional differences are unlikely to reflect adaptive convergence toward host GC content, since host genomes themselves vary only modestly (58% in *T. striata* vs. 53% in *T. suecica* and *T. chui*).

Regions of gene colinearity were apparent between TetV-1 and TetV-2 but were frequently interrupted by large-scale genomic rearrangements (**Fig. 1D**). The TetV-2 genome contained eight nearly identical copies (97.5–100% nucleotide identity) of a gene doublet encoding a bacterial transposase-related protein and an uncharacterized “transposase-associated” protein. These duplications likely contributed to the failure of short-read assembly. By contrast, TetV-1 contained only a single copy of this doublet. Phylogenetic analyses indicated that the duplications occurred after divergence of the TetV-1 and TetV-2 lineages (**Fig. S2**). Several TetV-2 doublets were positioned at the boundaries of colinear regions, suggesting a role in mediating homologous recombination and genome rearrangements.

Given their close relationship yet distinct host specificities, we propose that TetV-2 be assigned to the genus *Oceanusvirus* as the first representative of a new species, *O. lionense*, named after the “salin du Lion” site where it was first isolated.

### TetV-2 infection triggers reactivation of integrated PLVs

Analysis of the initial short-read assembly revealed five contigs (1,775–17,559 bp) matching endogenous Tsv elements (ETEs) previously identified in the T. striata LANL1001 genome, with >99% nucleotide identity across their entire length (**Fig. S3**) [23]. Short-read mapping against the LANL1001 reference uncovered ten additional regions with significant coverage, all but one overlapping known ETEs, including four fully spanned between terminal inverted repeats (TIRs; C2208, C1731, C0566, C2184). Two of these (C2208, C2184) were also supported by long reads from the TetV-2 lysate, indicating that their corresponding viral genomes were present in an independent infection experiment. No other LANL1001 regions were supported by long-read data.

Although the BG host strain used here differs from LANL1001, both belong to the same *T. striata* species group and share near-identical marker genes (100% identity over a 780 bp RBCL fragment; 99.8% identity over a 1,542 bp 18S fragment; **Fig. S4 and Supplementary Data**). Thus, the observed read recruitment cannot be attributed to host DNA release during lysis, which would yield random mapping patterns. Instead, the data indicate that the lysates contained additional Tsv-like viruses closely related to *T. striata* ETEs.

To reconstruct these genomes, we performed ETE-guided local assemblies for the four fully covered elements. Each yielded a single contig flanked by TIRs, corresponding to full-length Tsv-like genomes designated Tsv-S2a, Tsv-S2a.bis, Tsv-S2b, and Tsv-S3b (hereafter S2a, S2a.bis, S2b, and S3b; **Fig. 2**).

**Figure 2.**
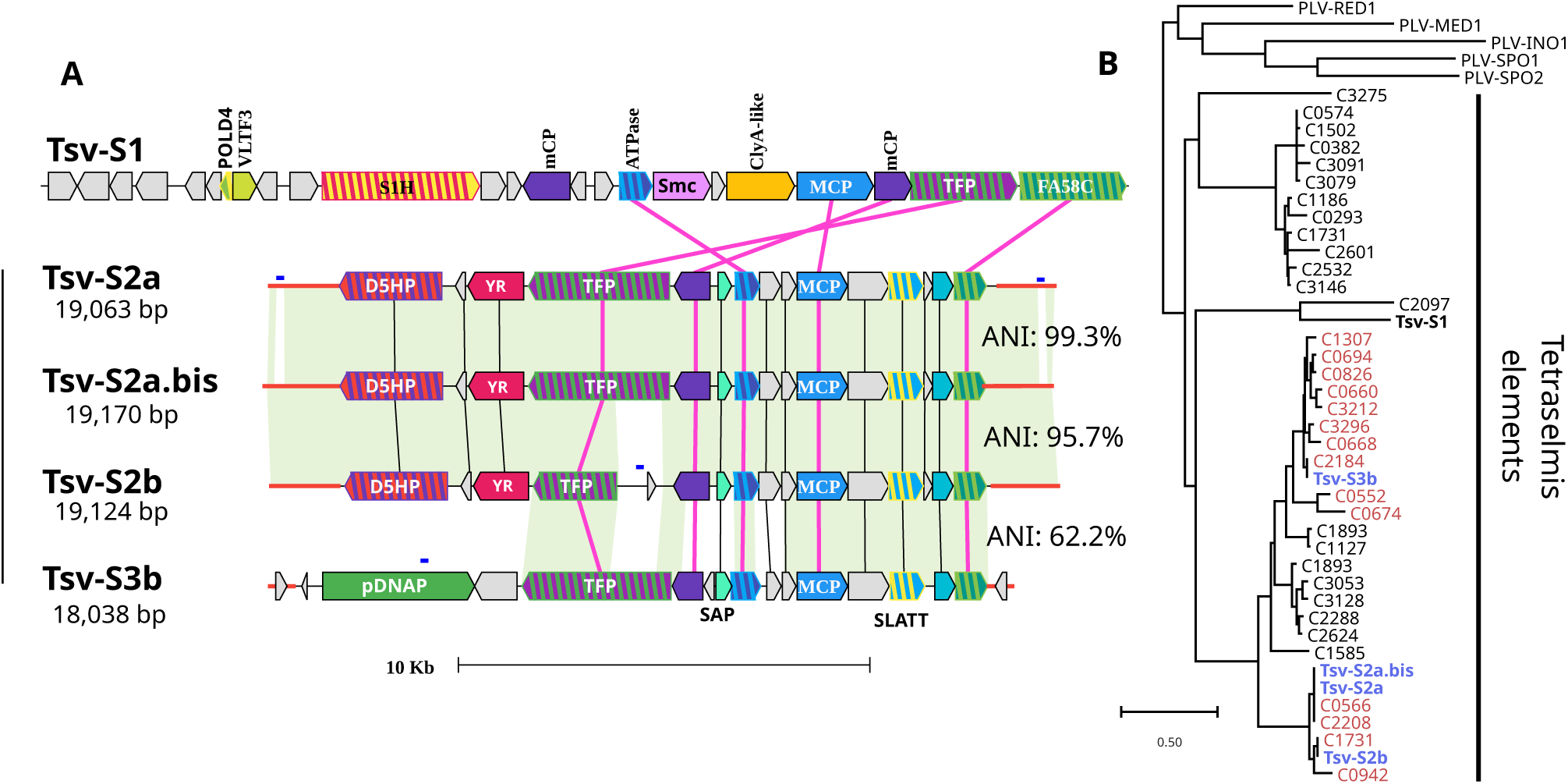
Tetraselmis striata Tsv genomes. (A) Genome maps of Tetraselmis striata viruses. Genes with annotated functions are shown as colored rectangles; hypothetical genes are shown in gray. Genes with significant protein-level similarity (BLASTP) are connected by lines, and regions with significant nucleotide similarity (BLASTN) are highlighted in green. Pink lines indicate genes conserved across all five viruses. Terminal inverted repeats are marked in red at the genome extremities. Average nucleotide identities (ANI) between genomes were calculated as the mean across BLASTN high-scoring pairs (HSPs). (B) Maximum-likelihood phylogenetic tree of the MCPs. Branch support values correspond to approximate likelihood-ratio test (aLRT) local supports (only values <80 are shown). PLV sequences used as outgroups (RED1, MED1, INO1, SPO1, and SPO2) were obtained from Yutin et al. (2015). Tsv-like elements detected by lysate sequencing are highlighted in red. Abbreviations: POLD4, DNA polymerase delta subunit 4; VLTF3, very late transcription factor 3; S1H, superfamily 1 helicase fused to an inactivated pDNAP domain; mCP, minor capsid protein; ATPase, DNA packaging ATPase; Smc, structural maintenance of chromosomes domain; Clya-like, ClyA-like α-pore-forming toxin domain; MCP, major capsid protein; TFP, tail fiber protein; FA58C, coagulation factor 5/8 C-terminal (discoidin) domain; D5HP, helicase–primase D5; SAP, SAP DNA-binding domain protein; AOx, PLN02976 amine oxidase domain; pDNAP, protein-primed family B DNA polymerase; SLATT_4, SMODS and SLOG-associating 2TM effector domain family 4; YR, tyrosine recombinase.

PCR assays with subtype-specific primers confirmed the presence of **S2a** and **S3b** sequences both in uninfected *Tetraselmis* BG host DNA and as encapsidated DNA in filtered lysates, consistent with the reactivation of endogenous elements. In contrast, **S2b** DNA was detected exclusively in lysates, suggesting that it was co-isolated with TetV-2 from the environmental sample rather than derived from the host genome. This interpretation was further supported by serial passages of purified TetV-2, which consistently yielded lysates containing S2a and S3b but never S2b, demonstrating that S2b is not generated through endogenous element reactivation. **S2a.bis** could not be assessed, as its sequence lacks unique regions suitable for primer design that would allow discrimination from other TsV variants.

Insufficient sequencing depth prevented assembly of the remaining ETEs, but mapping patterns suggested additional low-abundance, likely truncated, Tsv-like fragments. The four genomes (18,038–19,170 bp; **Table S1**) were substantially smaller than the free-living TsV-S1 genome (26,407 bp [22]). TIRs ranged from 583 bp (S3b) to 1,724 bp (S2b) and were absent from TsV-S1. The three S2x genomes (i.e., S2a, S2a.bis and S2b) terminated with monotonous “AT” dinucleotide repeats (**Fig. S5**), also present in their ETE guides, whereas S3b and its guide C2184 lacked such repeats. Each contig shared very high nucleotide identity with its ETE guide (ANI = 99.1–100%; **Table S2**). Notably, S2a.bis was slightly more similar to S2a (ANI = 99.3%) than to its guide C0566 (ANI = 99.1%), suggesting it does not derive from reactivation of a C0566 ortholog in BG. The S2b contig also differed from its guide, which carried a 5,441 bp tandem duplication absent from the viral genome (**Fig. S6**). By contrast, S3b was more divergent (ANI = 62.2–66.1% with the S2x group), with similarity limited to the distal two-thirds of the genome. No significant nucleotide similarity was detected between TsV-S1 and any contig by BLASTN, although BLASTP readily identified homologous proteins.

Phylogenetic analysis of the major capsid protein (MCP) confirmed these relationships (**Fig. 2B)**: the four novel Tsv-like viruses clustered more closely with each other than with TsV-S1, forming two clades—S2x and S3b. Short-read-covered LANL1001 ETEs (highlighted in red in **Fig. 2B**) also grouped into these two clades, indicating that ETE reactivation was not random but restricted to subsets of related elements, possibly responding to regulatory cues triggered by TetV-2 replication. We cannot exclude, however, that fewer Tsvs were reactivated and that lower/partial coverage of other ETEs reflects spurious read alignments due to sequencing errors.

Gene content analysis showed that S2a, S2a.bis, and S3b each encoded 16 predicted proteins, whereas S2b encoded 17. All four shared five conserved genes with TsV-S1: the major and minor capsid proteins, a DNA-packaging ATPase, a tail fiber protein (TFB), and a coagulation factor 5/8 C-terminal domain (FA58C)-like protein likely involved in carbohydrate binding [30]. The TFB gene in TsV-S2b was severely truncated compared to its homologs in the other PLVs. These conserved genes were clustered on one side of the genome, with clade-specific genes located on the opposite side. Unique regions were enriched in replication functions. TsV-S1 encoded a superfamily 1 helicase fused to an inactive protein-primed DNA polymerase domain (S1H) [31]; S3b encoded a protein-primed family B DNA polymerase (pDNAP) most similar to homologs in Polinton transposable elements [11]; and the S2x group encoded a D5 helicase–primase (D5HP) adjacent to a tyrosine recombinase, potentially mediating proviral integration and excision [32]. Beyond the core capsid and replication modules, the novel TsVs shared five additional genes absent from TsV-S1, including a SAP DNA-binding protein, a SLATT-domain protein related to predicted membrane-perforating toxins [33], and three uncharacterized proteins.

### Ultrastructural analysis

Transmission electron microscopy (TEM) provided direct evidence of viral infection within Tetraselmis cells and revealed the presence of two distinct types of virus-like particles (**Fig. S7**). Negative staining of cell lysates showed large icosahedral particles measuring 214–239 nm in diameter, consistent in size and morphology with *Tetraselmis* virus 1 (TetV-1; 226–257 nm) and therefore identified as *TetV*-2 virions. A second population of smaller icosahedral particles, 66–75 nm in diameter, corresponded to the size range previously reported for *Tetraselmis* virus N1 (TsV-N1; 49–73 nm). The coexistence of these two particle classes indicates concurrent production of a giant virus (*TetV*-2) and smaller TsV-like viruses.

To characterize the replication dynamics of *TetV*-2, thin-section TEM was performed on *T. striata* cells at 24, 48, and 72 hours post-inoculation (hpi). At 24 hpi, infected cells displayed extensive cytoplasmic reorganization, with large regions occupied by empty and filled *TetV*-2 capsids arising from membranous precursors (**Fig. S7 A1-C2**). These features closely resembled those described for *Tetraselmis* sp. infected with *TetV*-1 [34], including putative cytoplasmic viral factories with spatially organized capsid maturation, immature particles located centrally and mature virions peripherally. A minority population of smaller icosahedral particles with electron-dense cores was also observed, likely corresponding to TsV-like viruses replicating in parallel with *TetV*-2.

By 48 hpi, the number of *TetV*-2 particles had markedly declined, while TsV-like particles became predominant, frequently accumulating at the cell periphery near chloroplast membranes rather than in the central cytoplasm where *TetV*-2 factories had formed (**Fig. S7 D1-F2**). This shift in viral abundance suggests a temporal succession in which TsV replication becomes predominant after the peak of TetV-2 particle production.

At 72 h.p.i., TetV-2 particles were rarely detected, and the cytoplasm was almost entirely filled with TsV-like particles (**Fig. S7 G1-I2**). These smaller particles often formed crystalline arrays, a hallmark of advanced viral replication and accumulation. Collectively, these ultrastructural observations demonstrate that the polinton-like virus (PLV) genomes identified by metagenomic sequencing correspond to actively replicating viruses that co-occur with, but are distinct from, the giant *TetV*-2.

To verify encapsidation of the TsV-like viruses, we applied a size-fractionation and DNase-protection assay previously used to demonstrate the virion nature of Gezel-14T [35]. A fresh lysate was sequentially filtered through 0.45-µm and 0.1-µm membranes to remove intact cells and giant viruses, respectively. Aliquots of the 0.1-µm filtrate were boiled to release encapsidated DNA. The clarified (0.45-µm), filtered (0.1-µm), and boiled fractions were then treated with DNase to degrade unprotected DNA prior to PCR amplification (**Fig. S8**). PCR detection of specific marker genes revealed encapsidated *TetV*-2 DNA exclusively in the clarified fraction, whereas *TsV*-like markers were detected in both the clarified and filtered fractions. These results indicate that *TetV*-2 and *TsV*-like viruses are independently encapsidated, forming large (>0.1 µm) and small (<0.1 µm) virions, respectively.

### TsVs exhibit a virophage-like lifestyle

To evaluate whether TsVs act as virophages, we investigated their impact on the replication dynamics of the giant virus TetV-2. Host cultures were coinfected with a fixed concentration of TetV-2 (3.10^6^ ml^-1^) and varying ratios of TsV to TetV-2. Two distinct TsV cocktails derived from independent lysates were tested: one dominated by S2b (i.e., 99.4% S2b, 0.6% S3b; 0,0 % S2a but still detectable by qPCR [est. 8.2 copies per ml]) and the other dominated by S2a (99.4% S2a, 0.6% S3b; S2b undetectable). Cultures were incubated for 13 days, after which both host cells and viral particles were enumerated.

Control infections with TsVs alone yielded no detectable TsV progeny (**Fig. S9**) and did not increase host mortality (**Fig. 3A**). Instead, TsV abundance declined by more than 98% between inoculation and the end of the experiment, likely reflecting adsorption to host cell surfaces and virion degradation. This confirmed that TsVs cannot replicate autonomously in this host and are not directly harmful to the host. By contrast, coinfections with TetV-2 revealed a strong inhibitory effect of both TsV cocktails on giant virus production (**Fig. 3B**). Increasing the TsV/TetV-2 ratio in the inoculum caused a log-linear, dose-dependent reduction in TetV-2 yields. These findings indicate that TsVs require TetV-2 for propagation and, in turn, suppress TetV-2 replication, consistent with a virophage-like lifestyle. Nevertheless we did not observe a measurable reduction in host mortality in cultures co-infected with TetV-2 and TsVs; by day 13, all such cultures had visibly clarified.

**Figure 3.**
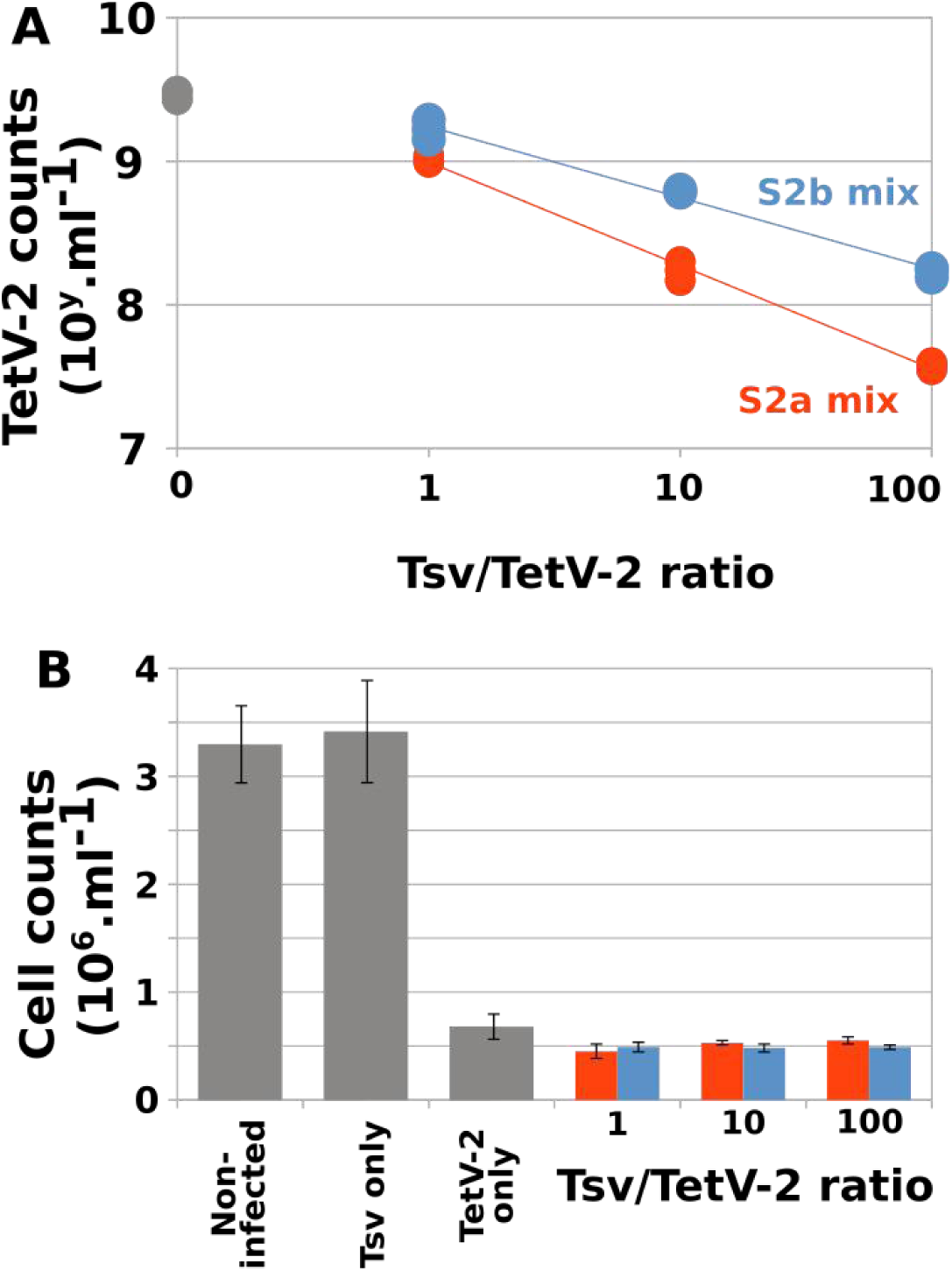
Virophage-associated life traits of TsV. (A) Abundance of TetV-2 particles at 13 days post infection (dpi) as a function of the initial TsV:TetV-2 inoculation ratio. Cultures were co-infected with TetV-2 (3 × 10⁶ particles mL⁻¹) and a TsV mixture dominated by strain TsV-S2a (red) or TsV-S2b (blue). The TetV-2–only control is shown in gray. Each point represents an independent biological replicate (n = 3), and lines depict the corresponding regression fits. **(B)** Concentration of viable Tetraselmis cells at 13 dpi. Gray bars indicate control cultures: uninfected, infected with the TsV-S2b–dominant mix only (10 × 10⁶ particles mL⁻¹ total TsV), or infected with TetV-2 alone. Red and blue bars correspond to co-infections with TetV-2 and the TsV-S2a– or TsV-S2b–dominant mixes, respectively. Bars represent mean values (n = 3), and error bars denote standard deviations.

### Replication cycle and latency phase

We investigated the replication dynamics of TetV-2 and reactivation of Tsv through a triplicate infection experiment, using cultures inoculated with purified TetV-2 particles. Samples were collected over an 8-day period post-inoculation. Host and viral abundances were monitored by flow cytometry, while extracellular encapsidated Tsv genome copies were quantified by qPCR.

The infection dynamics revealed that the latency phase of TetV-2 last between 21 and 24 hours (**Fig. 4**), which is longer than the reported 16-hours latency phase of TetV-1 [34]. After this period, TetV-2 abundance increased rapidly until ∼3 days post-infection (d.p.i.), after which it stabilized and remained constant until the end of the experiment. Detection of the two endogenous polinton-like viruses (Tsv-S2a and Tsv-S3b) was delayed relative to TetV-2: Tsv-S2a was first detected at 24 hours post-infection (h.p.i.), while measurable amounts of Tsv-S3b appeared only at 2 d.p.i. Unlike TetV-2, both PLVs continued to accumulate throughout the experiment without reaching a plateau. By the final sampling point, Tsv-S2a abundance exceeded that of TetV-2 by ∼9-fold and that of Tsv-S3b by ∼452-fold.

**Figure 4.**
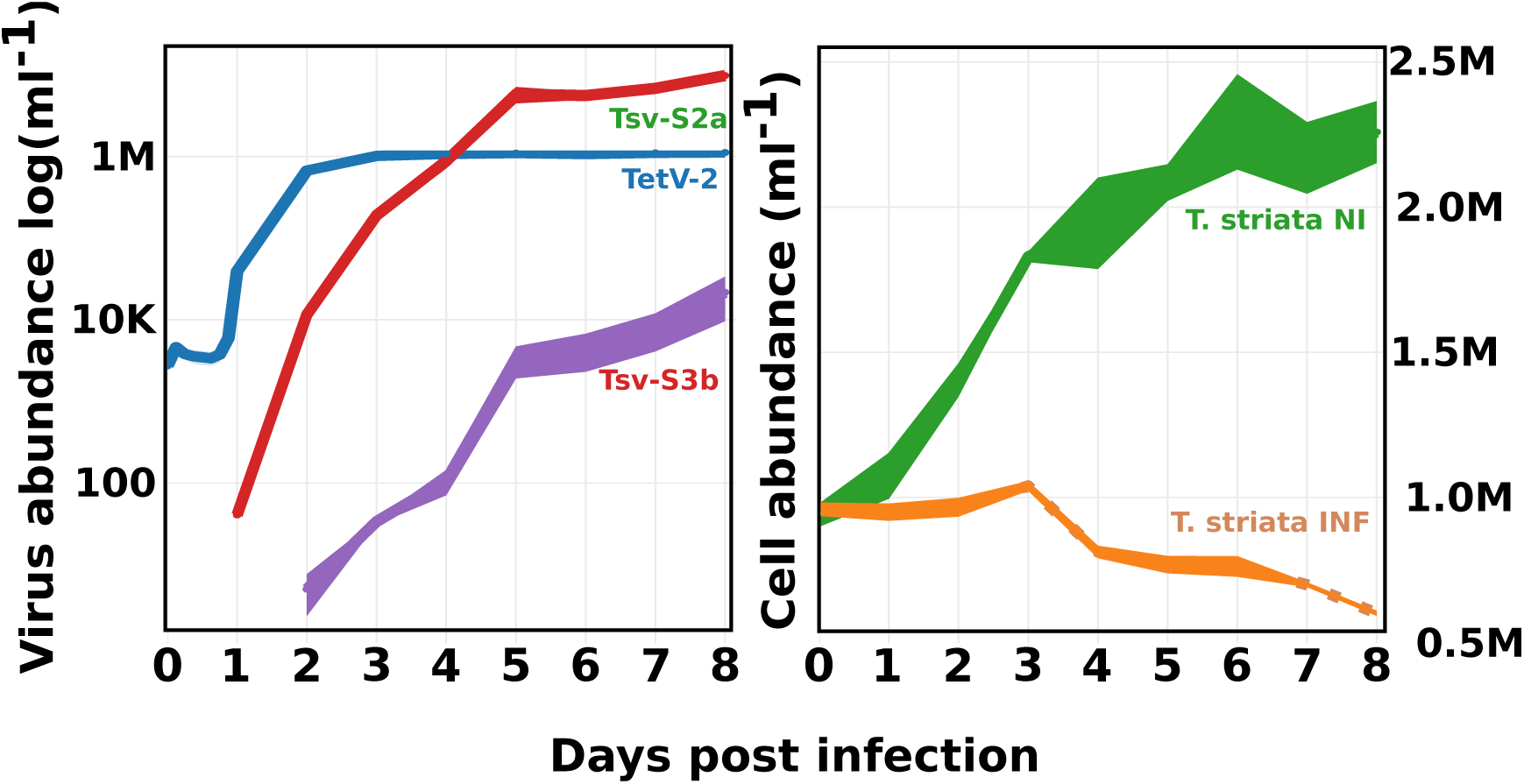
Infection dynamics of Tetraselmis by TetV-2. (A) Replication dynamics of TetV-2, TsV-S2a, and TsV-S3b following infection at a ratio of 30 TetV-2 particles per host cell. TetV-2 particle abundance was quantified by flow cytometry, while encapsidated DNA copy numbers of TsV-S2a and TsV-S3b were determined by qPCR. **(B)** Host cell survival over time, assessed by flow cytometry in infected and uninfected control cultures. Each point represents the mean of three biological replicates, and the shaded areas indicate the standard deviation.

These quantitative patterns are consistent with TEM observations. At 24 h.p.i., the cytoplasm of infected cells contained predominantly TetV-2 particles, with only a few Tsv particles visible, reflecting the initial burst of giant virus replication. By 72 h.p.i., however, giant viral particles were largely absent, and cells were densely filled with small Tsv particles. This transition in particle composition mirrors the dynamics observed in abundance profiles: the stabilization of TetV-2 levels after 3 d.p.i. corresponds to the progressive and sustained amplification of the virophages, which continued to accumulate until the end of the experiment.

### Host range of TetV-2

The host range of TetV-2 was assessed across seven *Tetraselmis* strains, including six *T. striata* isolates and the original *T. striata* BG strain from which TetV-2 was isolated (**Fig. S10**). Cultures were inoculated with purified TetV-2 and monitored after 13 d.p.i.. Cell concentrations and TetV-2 particle abundances were quantified by flow cytometry, while TsV DNA copy numbers were determined by qPCR.

The tested *Tetraselmis* strains displayed variable susceptibility to TetV-2, reflected in host mortality and viral productivity (**Fig. 5**). The highest mortality occurred in *T. striata* BG and RCC126, with cell densities reduced by 80–90% relative to uninfected controls at 13 d.p.i. In contrast, strains TetD, TetF, and LANL1001 displayed moderate reductions in cell abundance (7–36%), while *T. suecica* exhibited enhanced growth in inoculated cultures (+39%). The latter response likely reflects stimulation by material present in the inoculum, such as submicron cell debris not removed during the 0.45 µm prefiltration step.

**Figure 5.**
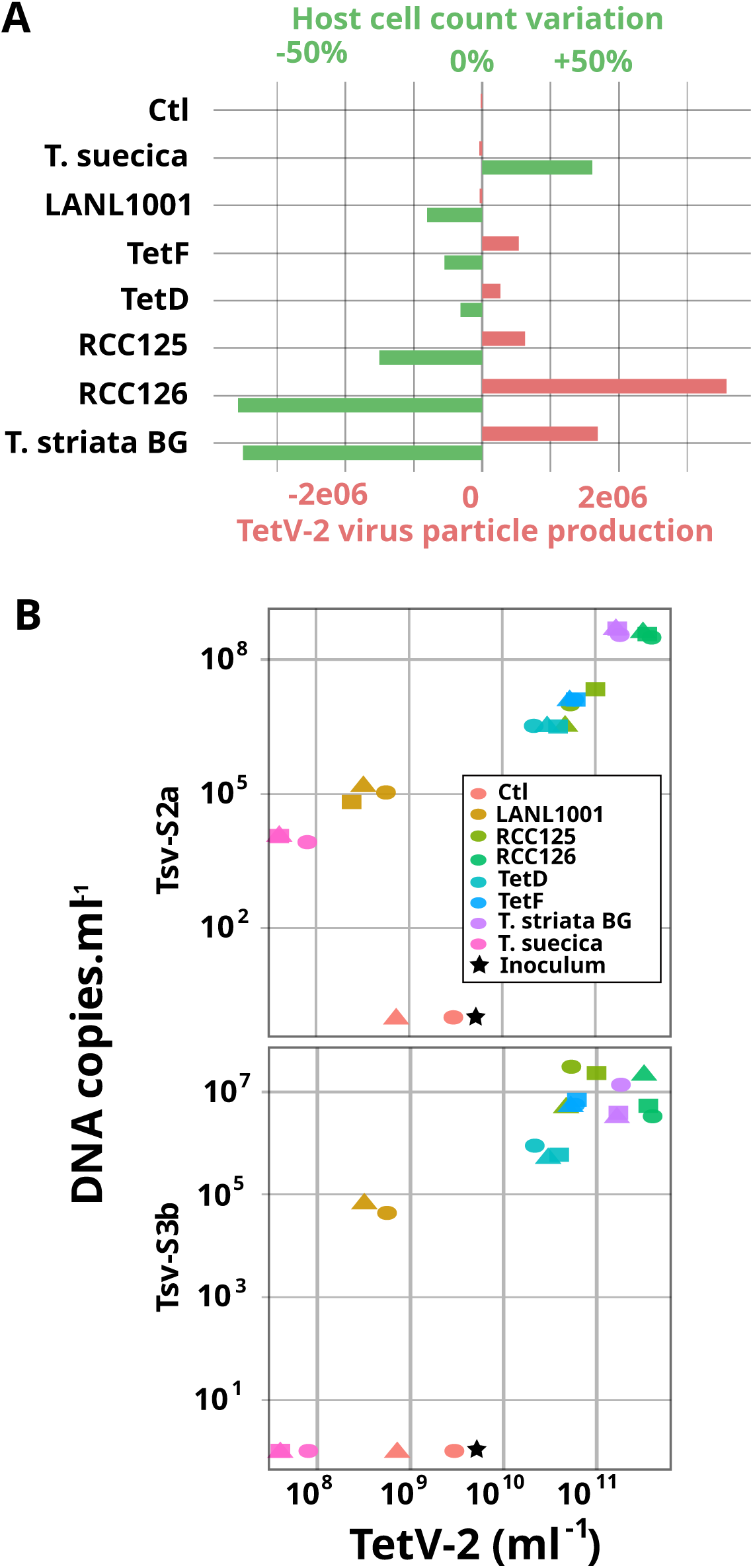
Host range of TetV-2 and productivity of Tsv. (A) Impact of TetV-2 infection on various Tetraselmis hosts after 13 days of incubation with a purified TetV-2 inoculum. Green bars indicate the variation in host cell abundance between infected and control (non-infected) cultures (mean, n = 3). Red bars represent the net increase in TetV-2 particle abundance relative to the number of particles initially inoculated (mean, n = 3). (B) Quantification of Tsv DNA copies produced by TetV-2–infected cultures as a function of TetV-2 net production. Each data point corresponds to an individual biological replicate.

Quantification of *TsV*-S2a revealed distinct, strain-dependent patterns. *TsV*-S2a DNA was consistently detected across all *Tetraselmis* strains, with copy numbers generally correlating with *TetV*-2 production. Notably, *TsV*-S2a was detected in infected LAN1001 cultures despite the absence of detectable *TetV*-2 replication, and low but reproducible *TsV*-S2a levels were also observed in *T. suecica*, which did not support *TetV*-2 propagation. *TsV*-S2a–specific sequences were present in the genomes of all *Tetraselmis* strains except RCC126, TetF, and *T. suecica* **(Table S3)**. These findings suggest that in these latter strains, small amounts of *TsV*-S2a introduced with the *TetV*-2 inoculum (below the qPCR detection limit) were nevertheless able to replicate within the algal host in the presence of the giant virus. Replication of *TsV*-S3b was detected in all *Tetraselmis* cultures except *T. suecica*, with copy numbers generally correlating with *TetV*-2 production to some extant. *TsV*-S3b sequences was detected in the same Tetraselmis strains which show *TsV*-S3b replication.

Together, these results reveal a complex landscape of Tetraselmis susceptibility to TetV-2 infection, shaped by both host-intrinsic factors and the presence of co-replicating PLVs. The observation that TsV-S2a can replicate in strains where TetV-2 replication is limited or undetectable suggests that PLVs may exploit transient windows of giant virus activity. The detection of TsV activity across diverse Tetraselmis strains, including those not permissive to TetV-2, indicates that PLVs may contribute to maintaining viral genetic diversity and influence infection outcomes at the population level. These interactions may attenuate the fitness of the giant virus, as observed in T. striata BG, or modulate infection dynamics in less susceptible strains. From an ecological perspective, such relationships could promote host population resilience by buffering against large-scale viral mortality events, thereby sustaining Tetraselmis bloom persistence.

## Discussion

We describe a natural coinfection system in *T. striata* in which a giant virus, *Oceanusvirus lionense* TetV-2, triggers the productive replication of multiple PLVs, one co-isolated from the same seawater sample (Tsv-S2b) and at least two elements integrated in the algal genome (Tsv-S2a, Tsv-S3b). Three lines of evidence indicate that one or more of these PLVs behave as virophages in this context: (i) the PLVs fail to propagate in the absence of TetV-2 and are not cytotoxic on their own; (ii) their production is temporally delayed relative to TetV-2 and then strongly predominates, consistent with helper-dependent amplification; and (iii) increasing PLV:TetV-2 input ratios reduces TetV-2 yields in a dose-dependent fashion. Together, these traits substantiate a parasitic interaction in which PLVs exploit the TetV-2 replication program and, in turn, diminish giant virus fitness. These results support the hypothesis that Polinton-like virophages can act as an adaptive antiviral defense system in Tetraselmis algae. However, because our current model system does not yet allow discrimination between the different *Tetraselmis*-derived PLVs produced during infection, it remains unclear whether one, several, or all of the Tsv elements characterized here possess virophage activity.

This system fills a significant gap in algal virology. Prior observations established that PLVs can exist as autonomous lytic viruses (e.g., TsV-N1/TvV-S1) [22,28] or as integrated elements in eukaryotic genomes [17,19,20,23] and are highly prevalent in aquatic systems [18,36]. Co-infection of giant virus and PLVs have been documented for two haptophyte algae [35,37,38]. PLV-like elements have also been found in association with entomopoxviruses infecting lepidopteran insects, establishing the first known link between PLVs and poxviruses [39]. Our findings add direct experimental support that, in *T. striata*, integrated PLVs can be reactivated by a giant virus and pursue a virophage-like lifestyle. Thus, *T. striata* hosts a continuum of PLV states, i.e., integrated, giant-virus-dependent, and free-living, within a single algal lineage, offering a powerful platform to interrogate transitions between mobile element and viral lifestyles.

TetV-2 diverges markedly from *O. kaneohense* TetV-1, having lost multiple metabolic and DNA-repair genes while expanding a transposase-associated gene doublet that coincides with local breaks in gene colinearity. These duplications may facilitate homologous recombination and genome rearrangement, potentially accelerating adaptation to host defenses or to parasitic PLVs. The pronounced GC-content divergence (TetV-2, 57.8%; TetV-1, 41.2%) relative to more modest host variation suggests long-term, lineage-specific mutational pressures, possibly reflecting distinct ecological niches or differential exposure to PLV parasitism. Recent work by Gajigan *et al.* (2025) [34] characterized infection of a *Tetraselmis* strain by TetV-1 through integrated transcriptomic and ultrastructural analyses. They reported extensive cytoplasmic reorganization and a well-defined viral factory, together with broad, temporally structured viral gene expression that included atypical metabolic functions linked to carbohydrate degradation and fermentation. Notably, no smaller virus-like particles or transcriptional signatures attributable to PLV-like elements were reported, although transcripts corresponding to integrated giant virus genes, including those encoding TetV-1–like major capsid proteins (MCPs), were observed. To probe this explicitly, we re-analyzed their published RNA-seq data by assembling transcripts and aligning them against an in-house database of integrated PLV sequences from Chlorodendrophyceae genomes [23]. Consistent with their observations, we found no evidence for the transcriptional reactivation of endogenous PLVs during TetV-1 infection. It remains possible that the genome of this *Tetraselmis* strain lacks integrated PLV elements, an unprecedented feature since all six *Chlorodendrophyceae* genomes sequenced to date harbor such integrations [23]. Alternatively, TetV-1 may not induce PLV reactivation under the experimental conditions reported, implying that only specific Oceanusvirus–PLV pairings can trigger such interactions. This interpretation would align with our observations for TetV-2, which reactivates only a subset of integrated PLVs in *T. striata*, suggesting a degree of molecular specificity between Oceanusvirus and PLV lineages.

All Tetraselmis PLVs share a conserved structural module, including major/minor capsid proteins (MCP/mCP), packaging ATPase, a putative tail fiber, and a FA58C-like carbohydrate-binding protein that colocalized on one genomic arm. In contrast, the replication module varies by clade (S2x vs S3b) and differs from the autonomous TsV-S1/TsV-N1 toolkit (e.g., S1 helicase fused to an inactive pDNAP). This partitioning supports a modular evolution scenario wherein structural genes are selectively maintained under capsid architecture constraints, whereas replication genes are replaced and modulate the PLV lifestyle autonomous or virophagous. Mechanistically, several lines of evidence point to how TetV-2 “awakens” PLVs. First, the terminal inverted repeats (TIRs) and the presence of tyrosine recombinase genes for S2x or protein-primed polymerase for S3b suggest that at least a subset of these elements can excise from the host genome and form replication-competent templates [39,40]. Second, the D5-like helicase–primase in S2x implies dependence on a helper-supplied replisome or on shared factory resources to initiate DNA synthesis [31]. Third, the temporal hierarchy, i.e., TetV-2 particles abundant at ∼24 h post-inoculation, followed by rising PLV titers from 24 h onward, argues that PLVs are triggered by factors that appear only after TetV-2 factory establishment, such as factory-localized transcriptional machinery, helper virus transcription factors, or nucleotide pools and membranes needed for virion assembly.

Although PLVs reduce TetV-2 yields in batch culture, they do not rescue *Tetraselmis* populations from collapse by day 13. In natural environments, however, even partial suppression of giant-virus productivity could alter algal bloom trajectories if it delays host lysis, reduces burst size, or increases infection heterogeneity. Strain-dependent PLV replication, including cases where PLV amplification occurs despite limited TetV-2 replication, suggests that PLVs can exploit transient or subthreshold giant-virus activity. Such dynamics could redistribute mortality across host subpopulations, buffer bloom collapse, and sustain viral genetic diversity by maintaining both giant virus and PLV lineages at intermediate frequencies.

All in all, our results show that numerous features are shared between the tripartite Tetraselmis–Oceanusvirus–PLV and Cafeteria–CroV–mavirus systems, despite the deep phylogenetic divergence of their hosts and associated viruses. In both protists, the host genome harbors a rich repertoire of integrated viral elements that can remain transcriptionally silent until reactivated by infection with a compatible giant virus. In Cafeteria burkhardae, the endogenous virophage mavirus is quiescent in uninfected cells but becomes transcriptionally and replicatively active upon CroV infection, a process requiring the giant virus late-phase transcriptional machinery and DNA replication. Inhibition of host or CroV protein synthesis abolishes this reactivation, confirming that provirophage induction depends on the functional state of the viral factory rather than on host stress responses [13]. The reactivated mavirus markedly suppresses CroV production in subsequent infection cycles, providing an altruistic, population-level antiviral effect. Genomic analyses of Cafeteria lineages further reveal a high diversity of endogenous virophages (EMALEs), which together occupy nearly 1– 2% of the nuclear genome [41]. These EMALEs cluster into distinct clades that differ in GC content, replication gene composition, and, critically, in their promoter motifs, implying molecular specificity toward different giant-virus partners. Only those EMALEs carrying CroV-like late promoters are transcriptionally reactivated upon CroV infection, illustrating a precise “compatibility” requirement between helper transcription factors and virophage regulatory sequences [15]. This architecture mirrors the modular organization of Tetraselmis PLVs, in which a conserved structural core is combined with divergent replication and integration modules, likely governing both their integration potential and their responsiveness to specific Oceanusvirus cues.

Thus, both systems embody a continuum of virophage states, i.e., integrated, giant-virus-dependent, and free-living, spanning parasitic and defensive roles. The giant-virus-specific induction of PLVs by TetV-2, together with the dose-dependent reduction of TetV-2 yields, closely parallels the Cafeteria model in both mechanism and ecological consequence, supporting the notion that Polinton-like virophages represent a widespread, evolutionarily conserved form of adaptive antiviral defense among unicellular eukaryotes.

## Materials and methods

### Algal Strains and Culture Conditions

Several *Tetraselmis* strains were used in this study, including *T. striata* BG, kindly provided by Gunnar Bratbak (University of Bergen, Norway); *T. striata* TetF and TetD, both isolated from a culture pond in Palavas-les-Flots, France, in September 2018; *T. striata* RCC125 and RCC126 obtained from the Roscoff Culture Collection (RCC, France); *T. striata* sp. LAN1001, and *T. suecica* CCMP904, kindly provided by Claire Sanders (Los Alamos National Laboratory, USA) and Sabine Stachowski Haberkorn (IFREMER, Nantes, France), respectively.

All strains were maintained in Conway medium (Ref. 33665256) at 20 °C under a 12:12 h light:dark photoperiod with an average irradiance of 1,500 lm m⁻² s⁻¹. Cultures were grown in 60 mL sterile plastic flasks and routinely subcultured every two weeks to maintain exponential growth.

#### Genomic DNA Extraction

For genomic DNA extraction used in quantitative PCR (qPCR) assays, 10 mL of each *Tetraselmis* culture were harvested by centrifugation at 1,500 × g for 10 min. Cell pellets were processed using the DNeasy PowerSoil Pro Kit (Qiagen) according to the manufacturer’s instructions.

### Virus isolation and diagnostic sequencing

A surface water sample was collected from brackish water at the Salin du Lion (43.45041°N, 5.22359°E; Vitrolles, France) on September 5, 2022. The sample was prefiltered through 0.45 µm membrane filters (supplier and reference number) to remove larger particles and eukaryotic cells. A 250 mL aliquot of the filtrate was amended with 10% (v/v) Conway medium and inoculated with 5 mL of an exponentially growing *Tetraselmis striata* GB culture. The co-culture was incubated for two weeks under standard growth conditions until host cell lysis was observed.

Following host clearance, the lysate was sequentially filtered through a 0.45 µm syringe filter and concentrated approximately 50-fold using a 100 kDa tangential flow filtration (TFF) system (Spectrum Laboratories). The concentrated viral fraction was treated with 40 U of Turbo DNase (Invitrogen) in 1× DNase buffer at room temperature to remove free nucleic acids. Viral nucleic acids were then extracted using the High Pure Viral Nucleic Acid Large Volume Kit (Roche) according to the manufacturer’s protocol. The purified DNA was used to construct sequencing libraries with the Nextera XT DNA Library Preparation Kit (Illumina) and sequenced on an Illumina MiSeq platform (2 × 250 bp paired-end) to detect and characterize viral genomes associated with the culture collapse.

A clonal *Tetraselmis* virus isolate, designated TetV-2, was obtained after three successive rounds of serial dilution to extinction. The PLV Tsv-S2b was co-isolated from the same environmental sample and was present in the initial lysate, whereas Tsv-S2a and Tsv-S3b were identified as endogenous elements integrated within the nuclear genome of *T. striata*. A PLV cocktail derived from the initial lysate, containing all three PLVs (Tsv-S2a, Tsv-S2b, and Tsv-S3b) in different proportions, was maintained by co-cultivation with TetV-2 and prepared by filtration of lysate through a 0.1 µm-pore membrane. A second PLV cocktail, derived from the clonal TetV-2 isolate, contained only the reactivated PLVs Tsv-S2a and Tsv-S3b; this preparation is produced on demand by inoculating *T. striata* cultures with purified TetV-2 and filtration of the resulting lysate through a 0.1 µm-pore membrane.

Inoculation with purified TetV-2: a 0.45 µm–filtered lysate is first quantified for TetV-2 particle abundance by flow cytometry. An appropriate volume of lysate (typically 50– 500 µL) is then diluted 1:100 in fresh Conway medium to achieve the desired virus-to-cell ratio. The resulting suspension is filtered through a 0.22 µm–pore membrane (60300, Pall Corporation), allowing the majority of PLVs to pass through while retaining TetV-2 particles on the filter surface. The TetV-2–containing filter is immediately introduced into a fresh Tetraselmis culture to initiate infection. qPCR assays performed on DNA extracted from the filter consistently confirmed the absence of detectable PLV sequences, although it is likely that trace amounts of PLV particles remained associated with the membrane.

### TetV-2 Production and Preparation of Viral DNA for Nanopore Sequencing

An exponentially growing *Tetraselmis striata* GB culture (600 mL) was infected with 200 µL of a clonal TetV-2 lysate and monitored daily until complete host lysis was observed. Following lysis, the culture was clarified by centrifugation at 1,500 × g for 10 min at room temperature, and the supernatant was filtered through a 0.45 µm membrane filter (supplier and reference). The clarified lysate was treated with 240 U of Turbo DNase (AM1907, Thermo Fisher Scientific) in 1× Turbo DNase buffer and 430 U of RNase A (supplier and reference) for 1 h at 37 °C to remove free nucleic acids.

Viral particles were subsequently precipitated by adding polyethylene glycol (PEG) 8000 to a final concentration of 10% (w/v) and NaCl to 0.5 M, followed by gentle agitation overnight at 4 °C. The precipitated viruses were recovered by centrifugation at 8,000 × g for 20 min at 4 °C, and the resulting pellets were resuspended in 600 µL of 5 mM MgSO₄.

High-molecular-weight (HMW) viral DNA was extracted using the Monarch HMW DNA Extraction Kit for Tissue (New England Biolabs), following the manufacturer’s “Protocol for High Molecular Weight DNA (HMW DNA) Extraction from Phage.” DNA integrity was evaluated using a High Sensitivity DNA ScreenTape assay (Agilent 4200 TapeStation), and concentration was quantified with the Qubit dsDNA High Sensitivity Assay Kit (Thermo Fisher Scientific). The extracted HMW DNA was sequenced on an Oxford Nanopore GridION device using the Ligation Sequencing Kit (SQK-LSK109) and R9.4.1 flow cells. Basecalling was performed with the high-accuracy (HAC) model, yielding a total of 12.3 Gb of long-read data.

### Sequence analysis

Illumina short-read data were processed as follows: reads were trimmed and filtered to remove low-quality sequences using FastP [42], then assembled with MEGAHIT [43] using default parameters. Oxford Nanopore Technologies (ONT) long-read data were filtered, assembled, and curated following the Trycycler workflow [44]. Gene prediction was performed with Prodigal. Phylogenetic trees were inferred using IQ-TREE [45], and sequence similarity searches against the NCBI nonredundant (NR) protein database were conducted using MMseqs2 [46].

### Verification of Tsv virion encapsidation

The 0.45 µm–filtered viral lysate was divided into three fractions. The first fraction was left untreated (clarified lysate). The second fraction was filtered twice through 0.1 µm syringe filters (FL4537, Pall Corporation) to remove giant virus particles (filtered lysate). Aliquots of the 0.1 µm filtrate were then boiled for 10 min at 95 °C to disrupt viral capsids (boiled filtered lysate).

All fractions were subsequently treated with DNase to degrade free or unprotected DNA. Each 50 µL reaction contained the sample, 1× DNase buffer, and 6 U of Turbo DNase (AM1907, Thermo Fisher Scientific), and was incubated for 30 min at 37 °C. DNase was inactivated according to the manufacturer’s instructions.

DNA extracted from each fraction was analyzed by qPCR using specific primers targeting TetV-2 and the three PLVs (Tsv-S2a, Tsv-S2b, and Tsv-S3b). The detection of DNase-resistant PLV genomes was interpreted as evidence of encapsidated virophage particles.

### Tetraselmis strain sensitivity to Oceanusvirus

Exponentially growing cultures of each *Tetraselmis* strain (n = 7) were quantified by flow cytometry, adjusted to 1 × 10⁶ cells mL⁻¹, and distributed into 6-well plates (5 mL per well) in triplicate for both infected and uninfected control conditions. Filters containing purified TetV-2 were prepared on the day of the experiment and transferred directly into wells containing *Tetraselmis* cultures. Uninfected control cultures were maintained under identical conditions using filters pre-rinsed with 20 mL of Conway medium. Infections were monitored daily by light microscopy until complete lysis was observed in *T. striata* BG triplicates infected with purified TetV-2.

At days 0 and 13 post-infection, 200 µL from each well were sampled and filtered through a 0.45 µm filter plate (MAHVN4510, Millipore). Filtrates were treated with 6 U of Turbo DNase in 1× DNase buffer (AM1907, Thermo Fisher Scientific) for 30 min at 37 °C to degrade free DNA. Subsequently, 200 µL of Qiagen lysis buffer AL were added, and the mixtures were divided into 200 µL aliquots and stored at –80 °C until nucleic acid extraction for virophage quantification by qPCR. Nucleic acid extraction was performed using the QIAamp DNA Mini Kit (ref: xxx, Qiagen) according to the manufacturer’s instructions. All extractions were performed on a QIAcube automated workstation (Qiagen) to ensure standardization of the procedure and to minimize handling variability.

To quantify TetV-2 and host cell concentrations at day 13, two additional 200 µL aliquots per well were collected, preserved with glutaraldehyde (0.25% v/v final concentration) and Pluronic F-68 (0.01% v/v final concentration), flash-frozen in liquid nitrogen, and stored at –80 °C until analysis by flow cytometry.

### TetV-2 infection dynamics in T. striata BG

Triplicate *Tetraselmis striata* BG cultures (40 mL each) in exponential growth phase (1 × 10⁶ cells mL⁻¹) were inoculated with 0.22 µm filters containing purified TetV-2, corresponding to an estimated virus-to-host ratio of approximately 30:1. Uninfected control cultures were established in triplicate using the same procedure, but with TetV-2–free filters pre-rinsed with 20 mL of Conway medium.

Following inoculation, subsamples were collected periodically for up to 8 days post-infection. Duplicate 200 µL aliquots were preserved with glutaraldehyde (0.5% final concentration) and Pluronic F-68 (0.01% final concentration), flash-frozen in liquid nitrogen, and stored at –80 °C until flow cytometric quantification of TetV-2 particles and host cells.

For virophage nucleic acid extraction and quantification by qPCR, samples were processed as described for the above *Tetraselmis* strain sensitivity assay.

### Flow cytometer count

*Tetraselmis* cells and TetV-2 virus-like particles (VLPs) were enumerated using a CytoFLEX S flow cytometer (Beckman Coulter). Fixed samples were thawed in the dark at room temperature prior to analysis. Absolute quantification of both algal cells and TetV-2 VLPs was performed using BD Trucount™ Tubes (663028, BD Biosciences). Blanks consisting of Conway medium or pretreated 1× TE buffer were systematically analyzed in parallel with the samples.

For TetV-2 particle quantification, samples were diluted 20-fold in 500 µL of 0.22 µm– filtered 1× TE buffer (10530028-3, Bio-World), stained with 10 µL of a 1000× SYBR Green I solution (Thermo Fisher Scientific), vortexed for 15 s, incubated in the dark for 10 min at 80 °C, and cooled for 5 min at room temperature before analysis. To achieve an optimal event rate (200–800 events s⁻¹) and minimize electronic coincidence, additional dilution in 0.22 µm–filtered 1× TE buffer was performed when necessary. Data acquisition was triggered on green fluorescence, and VLPs were discriminated based on violet side scatter (VSSC) versus green fluorescence, following the approach described by Zhao *et al.* [47]. Data were collected in logarithmic mode using CytExpert software (v2.3.0.84, Beckman Coulter).

For *Tetraselmis* cell enumeration, algal cells were identified using side scatter (SSC) and chlorophyll autofluorescence (PC5.5) signals.

### Real time PCR

qPCR assays were used to quantify virophages and TetV-2 DNA in the different samples. Reactions were performed using the QuantiNova SYBR Green PCR Kit (208054, Qiagen) according to the manufacturer’s instructions, with primers at a final concentration of 0.5 µM. Four primer sets were used to specifically amplify virophages Tsv-S2a, Tsv-S2b, Tsv-S3b and TetV-2 **(Table S4)**. Thermal cycling conditions were optimized for each primer set. For Tsv-S2a, the program consisted of an initial denaturation at 95 °C for 2 min, followed by 40 cycles of 95 °C for 5 s, 55 °C for 30 s, and 72 °C for 30 s, ending with a melt curve analysis. For Tsv-S2b and Tsv-S3b, the program consisted of an initial denaturation at 95 °C for 2 min, followed by 40 cycles of 95 °C for 5 s and 60 °C for 15 s, also followed by a melt curve analysis.

Standard curves were generated from purified amplicons. PCR products were cleaned with the NucleoMag® kit for clean-up and size selection of NGS library preparation reactions (Macherey-Nagel) and quantified using a DNA HS Qubit™ Fluorometer (Q32854, Thermo Fisher Scientific). The Tsv-S2a primers target the TIR regions of the genome, which are present in duplicate. Consequently, the qPCR-estimated number of DNA targets for this PLV was divided by two to obtain the corresponding number of genome copies.

### Transmission Electron Microscopy Negative Staining

Samples fixed with glutaraldehyde 2.5% final were adsorbed directly onto glow-discharged formvar carbon films on 400 mesh nickel grids (FCF400-Ni, EMS) for 10 min at +4°C. Grids were stained for 10 s with 1% molybdate solution in filtered water at room temperature. Electron micrographs were obtained on a Tecnai G2 transmission electron microscope (Thermo-Fischer FEI) operated at 200 keV equipped with a 4096 × 4096 pixel resolution Eagle camera (FEI).

### Transmission Electron Microscopy of ultra-thin sections

Algae were fixed with 2.5 % glutaraldehyde in 0.1 M sodium cacodylate buffer. Resin embedding was microwave-assisted with a PELCO BiowavePro+. Samples were washed with a mixture of 0.2 M saccharose/0.1 M sodium cacodylate and post-fixed with 1% OsO4 diluted in 0.2 M potassium hexa-cyanoferrate (III)/0.1 M sodium cacodylate buffer. After being washed with distilled water, samples were gradually dehydrated by successive baths with 30% to 100% ethanol. Substitution with Epon resin was achieved by incubations with 25% to 100% Epon resin. After final ultracentrifugation at 2000 rpm for 2 minutes in order to get a low-density algae pellet, resin with algae was heat-cured for 72 h at 60 °C. All solutions used above were 0.2-μm filtered. Ultrathin 100 nm sections were cut using a UC7 ultramicrotome (Leica Microsystems) and placed on HR25 300 Mesh Copper/Rhodium grids (TAAB). Sections were contrasted with uranyl acetate and lead citrate according to Reynolds’s method [48]. Electron micrographs were obtained on a Tecnai G2 transmission electron microscope (Thermo-Fischer/FEI) operated at 200 keV equipped with a 4096 × 4096 pixels resolution Eagle camera (FEI).

## Supporting information

Supplementary Tables and Figures combined

## REFERENCES

1. Breitbart M. Marine Viruses: Truth or Dare. Annu Rev Mar Sci. 2012;4: 425–448. doi:10.1146/annurev-marine-120709-142805

2. Suttle CA. Marine viruses — major players in the global ecosystem. Nature Reviews Microbiology. 2007;5: 801–812. doi:10.1038/nrmicro1750

3. Fuhrman JA. Marine viruses and their biogeochemical and ecological effects. Nature. 1999;399: 541–548. doi:10.1038/21119

4. Hevroni G, Vincent F, Ku C, Sheyn U, Vardi A. Daily turnover of active giant virus infection during algal blooms revealed by single-cell transcriptomics. Sci Adv. 2023;9: eadf7971. doi:10.1126/sciadv.adf7971

5. Vardi A, Van Mooy BAS, Fredricks HF, Popendorf KJ, Ossolinski JE, Haramaty L, et al. Viral Glycosphingolipids Induce Lytic Infection and Cell Death in Marine Phytoplankton. Science. 2009;326: 861–865. doi:10.1126/science.1177322

6. Bratbak G, Egge J, Heldal M. Viral mortality of the marine alga Emiliania huxleyi (Haptophyceae) and termination of algal blooms. Mar Ecol Prog Ser. 1993;93: 39–48. doi:10.3354/meps093039

7. Forterre P, Gaïa M. Giant viruses and the origin of modern eukaryotes. Current Opinion in Microbiology. 2016;31: 44–49. doi:10.1016/j.mib.2016.02.001

8. Forterre P, Gaia M. Les virus géants et l’origine des ARN polymérases des eucaryotes. Med Sci (Paris). 2021;37: 230–233. doi:10.1051/medsci/2021007

9. Duponchel S, Fischer MG. Viva lavidaviruses! Five features of virophages that parasitize giant DNA viruses. Condit RC, editor. PLoS Pathog. 2019;15: e1007592. doi:10.1371/journal.ppat.1007592

10. La Scola B, Desnues C, Pagnier I, Robert C, Barrassi L, Fournous G, et al. The virophage as a unique parasite of the giant mimivirus. Nature. 2008;455: 100–104. doi:10.1038/nature07218

11. Koonin EV, Fischer MG, Kuhn JH, Krupovic M. The polinton-like supergroup of viruses: evolution, molecular biology, and taxonomy. Microbiol Mol Biol Rev. 2024; e0008623. doi:10.1128/mmbr.00086-23

12. Roux S, Fischer MG, Hackl T, Katz LA, Schulz F, Yutin N. Updated Virophage Taxonomy and Distinction from Polinton-like Viruses. Biomolecules. 2023;13: 204. doi:10.3390/biom13020204

13. Fischer MG, Hackl T. Host genome integration and giant virus-induced reactivation of the virophage mavirus. Nature. 2016;540: 288–291. doi:10.1038/nature20593

14. Fischer MG, Suttle CA. A Virophage at the Origin of Large DNA Transposons. Science. 2011;332: 231–234. doi:10.1126/science.1199412

15. Koslová A, Hackl T, Bade F, Sanchez Kasikovic A, Barenhoff K, Schimm F, et al. Endogenous virophages are active and mitigate giant virus infection in the marine protist *Cafeteria burkhardae*. Proc Natl Acad Sci USA. 2024;121: e2314606121. doi:10.1073/pnas.2314606121

16. Kapitonov VV, Jurka J. Self-synthesizing DNA transposons in eukaryotes. Proc Natl Acad Sci USA. 2006;103: 4540–4545. doi:10.1073/pnas.0600833103

17. Bellas C, Hackl T, Plakolb M-S, Koslová A, Fischer MG, Sommaruga R. Large-scale invasion of unicellular eukaryotic genomes by integrating DNA viruses. Proc Natl Acad Sci USA. 2023;120: e2300465120. doi:10.1073/pnas.2300465120

18. Bellas CM, Sommaruga R. Polinton-like viruses are abundant in aquatic ecosystems. Microbiome. 2021;9: 13. doi:10.1186/s40168-020-00956-0

19. Stephens D, Faghihi Z, Moniruzzaman M. Widespread occurrence and diverse origins of polintoviruses influence lineage-specific genome dynamics in stony corals. Virus Evolution. 2024;10: veae039. doi:10.1093/ve/veae039

20. Bulzu P-A, Henriques Vieira H, Ghai R. Lineage-specific expansions of polinton-like viruses in photosynthetic cryptophytes. Microbiome. 2025;13: 154. doi:10.1186/s40168-025-02148-0

21. Gyaltshen Y, Rozenberg A, Paasch A, Burns JA, Warring S, Larson RT, et al. Long-Read– Based Genome Assembly Reveals Numerous Endogenous Viral Elements in the Green Algal Bacterivore *Cymbomonas tetramitiformis*. Katz L, editor. Genome Biology and Evolution. 2023;15: evad194. doi:10.1093/gbe/evad194

22. Pagarete A, Grébert T, Stepanova O, Sandaa R-A, Bratbak G. Tsv-N1: A Novel DNA Algal Virus that Infects Tetraselmis striata. Viruses. 2015;7: 3937–3953. doi:10.3390/v7072806

23. Chase EE, Desnues C, Blanc G. Integrated viral elements suggest the dual lifestyle of *Tetraselmis* spp. Polinton-like viruses. Virus Evolution. 2022;8: veac068. doi:10.1093/ve/veac068

24. Fistarol GO, Coutinho FH, Moreira APB, Venas T, Cánovas A, De Paula SEM, et al. Environmental and Sanitary Conditions of Guanabara Bay, Rio de Janeiro. Front Microbiol. 2015;6. doi:10.3389/fmicb.2015.01232

25. Jones JB, Rhodes LL. Suffocation of pilchards (*Sardinops sagax*) by a green microalgal bloom in Wellington Harbour, New Zealand. New Zealand Journal of Marine and Freshwater Research. 1994;28: 379–383. doi:10.1080/00288330.1994.9516627

26. Pizarro M. An Unusual and Massive Bloom of Tetraselmis Sp. in the Valparaiso Bay, Chile. OFOAJ. 2018;7. doi:10.19080/OFOAJ.2018.07.555717

27. Schvarcz CR, Steward GF. A giant virus infecting green algae encodes key fermentation genes. Virology. 2018;518: 423–433. doi:10.1016/j.virol.2018.03.010

28. Sizov DV, Polischuk VP. Cultivation, purification and crystallization of virus of green algae Tetraselmis viridis. Biopolym Cell. 2006;22: 243–245. doi:10.7124/bc.000737

29. Aylward FO, Abrahão JS, Brussaard CPD, Fischer MG, Moniruzzaman M, Ogata H, et al. Taxonomic update for giant viruses in the order Imitervirales (phylum Nucleocytoviricota). Arch Virol. 2023;168: 283. doi:10.1007/s00705-023-05906-3

30. Gilbert GE, Baleja JD. Membrane-binding peptide from the C2 domain of factor VIII forms an amphipathic structure as determined by NMR spectroscopy. Biochemistry. 1995;34: 3022–3031. doi:10.1021/bi00009a033

31. Krupovic M, Yutin N, Koonin EV. Fusion of a superfamily 1 helicase and an inactivated DNA polymerase is a signature of common evolutionary history of Polintons, polinton-like viruses, Tlr1 transposons and transpovirons. Virus Evol. 2016;2: vew019. doi:10.1093/ve/vew019

32. Barth ZK, Hicklin I, Thézé J, Takatsuka J, Nakai M, Herniou EA, et al. Genomic analysis of hyperparasitic viruses associated with entomopoxviruses. Virus Evolution. 2024;10: veae051. doi:10.1093/ve/veae051

33. Burroughs AM, Zhang D, Schäffer DE, Iyer LM, Aravind L. Comparative genomic analyses reveal a vast, novel network of nucleotide-centric systems in biological conflicts, immunity and signaling. Nucleic Acids Res. 2015;43: 10633–10654. doi:10.1093/nar/gkv1267

34. Gajigan AP, Schvarcz CR, Conaco C, Edwards KF, Steward GF. Ultrastructural and transcriptional changes during a giant virus infection of a green alga. npj Viruses. 2025;3: 47. doi:10.1038/s44298-025-00128-7

35. Roitman S, Rozenberg A, Lavy T, Brussaard CPD, Kleifeld O, Béjà O. Isolation and infection cycle of a polinton-like virus virophage in an abundant marine alga. Nat Microbiol. 2023;8: 332–346. doi:10.1038/s41564-022-01305-7

36. Piedade GJ, Schön ME, Lood C, Fofanov MV, Wesdorp EM, Biggs TEG, et al. Seasonal dynamics and diversity of Antarctic marine viruses reveal a novel viral seascape. Nat Commun. 2024;15: 9192. doi:10.1038/s41467-024-53317-y

37. Santini S, Jeudy S, Bartoli J, Poirot O, Lescot M, Abergel C, et al. Genome of Phaeocystis globosa virus PgV-16T highlights the common ancestry of the largest known DNA viruses infecting eukaryotes. PNAS. 2013;110: 10800–10805. doi:10.1073/pnas.1303251110

38. Stough JMA, Yutin N, Chaban YV, Moniruzzaman M, Gann ER, Pound HL, et al. Genome and Environmental Activity of a Chrysochromulina parva Virus and Its Virophages. Front Microbiol. 2019;10: 703. doi:10.3389/fmicb.2019.00703

39. Barth ZK, Hicklin I, Thézé J, Takatsuka J, Nakai M, Herniou EA, et al. Genomic analysis of hyperparasitic viruses associated with entomopoxviruses. Virus Evolution. 2024;10: veae051. doi:10.1093/ve/veae051

40. Rajeev L, Malanowska K, Gardner JF. Challenging a Paradigm: the Role of DNA Homology in Tyrosine Recombinase Reactions. Microbiol Mol Biol Rev. 2009;73: 300–309. doi:10.1128/MMBR.00038-08

41. Hackl T, Duponchel S, Barenhoff K, Weinmann A, Fischer MG. Virophages and retrotransposons colonize the genomes of a heterotrophic flagellate. eLife. 2021;10: e72674. doi:10.7554/eLife.72674

42. Chen S, Zhou Y, Chen Y, Gu J. fastp: an ultra-fast all-in-one FASTQ preprocessor. Bioinformatics. 2018;34: i884–i890. doi:10.1093/bioinformatics/bty560

43. Li D, Liu C-M, Luo R, Sadakane K, Lam T-W. MEGAHIT: an ultra-fast single-node solution for large and complex metagenomics assembly via succinct *de Bruijn* graph. Bioinformatics. 2015;31: 1674–1676. doi:10.1093/bioinformatics/btv033

44. Wick RR, Judd LM, Cerdeira LT, Hawkey J, Méric G, Vezina B, et al. Trycycler: consensus long-read assemblies for bacterial genomes. Genome Biol. 2021;22: 266. doi:10.1186/s13059-021-02483-z

45. Nguyen L-T, Schmidt HA, von Haeseler A, Minh BQ. IQ-TREE: a fast and effective stochastic algorithm for estimating maximum-likelihood phylogenies. Mol Biol Evol. 2015;32: 268–274. doi:10.1093/molbev/msu300

46. Steinegger M, Söding J. MMseqs2 enables sensitive protein sequence searching for the analysis of massive data sets. Nat Biotechnol. 2017;35: 1026–1028. doi:10.1038/nbt.3988

47. Zhao Y, Zhao Y, Zheng S, Zhao L, Zhang W, Xiao T, et al. Enhanced resolution of marine viruses with violet side scatter. Cytometry Pt A. 2023;103: 260–268. doi:10.1002/cyto.a.24674

48. Reynolds ES. The use of lead citrate at high pH as an electron-opaque stain in electron microscopy. The Journal of Cell Biology. 1963;17: 208–212. doi:10.1083/jcb.17.1.208

