## Supplementary Tables and Figures combined for "A giant virus awakens polinton-like virophages in the green alga *Tetraselmis*, revealing an inducible antiviral defense system"

Table S1 : Features of Tetraselmis viruses

| Contig | Length (bp) | TIR length (bp) | Genome conformation | Number of protein genes | Number of tRNA genes |
|---|---|---|---|---|---|
| TetV-2 | 660 680 | None | Circular | 640 | 4 |
| Tsv-S2a | 19 063 | 1632 | Linear | 17 | 0 |
| Tsv-S2a.bis | 19 170 | 1711 | Linear | 17 | 0 |
| Tsv-S2b | 19 124 | 1724 | Linear | 18 | 0 |
| Tsv-S3b | 18 038 | 583 | Linear | 17 | 0 |

Table S2 : Average nucleotide identity between Tetraselmis viruses and their ETE guides

| Average Nucleotide Identity (ANI) | C2208 (S2a) | C1731 (S2b) | C2184 (S3b) | C0566 (S2a.bis) | Tsv-S2a | Tsv-S2a.bis | Tsv-S2b |
|---|---|---|---|---|---|---|---|
| C1731 (S2b) | 94,58 % | | | | | | |
| C2184 (S3b) | 65,31 % | 62,03 % | | | | | |
| C0566 (S2a.bis) | 98,53 % | 96,05 % | 65,81 % | | | | |
| Tsv-S2a | 99,98 % | 94,62 % | 65,37 % | 98,52 % | | | |
| Tsv-S2a.bis | 99,13 % | 95,58 % | 65,56 % | 99,05 % | 99,26 % | | |
| Tsv-S2b | 94,78 % | 99,81 % | 62,06 % | 96,01 % | 94,79 % | 95,72 % | |
| Tsv-S3b | 65,85 % | 62,18 % | 99,43 % | 65,96 % | 65,92 % | 66,11 % | 62,20 % |

Table S3 : Presence of Tsv target sequences in *Tetraselmis* genomic DNA.

| Strain | Tsv-S2a | Tsv-S2b | Tsv-S3b | PCR control | |
|---|---|---|---|---|---|
| | | | | 18S | EF1a |
| *T. striata* BG | ✔ | | ✔ | ✔ | ✔ |
| RCC126 | | | ✔ | ✔ | ✔ |
| RCC125 | ✔ | | ✔ | ✔ | ✔ |
| Tet D | ✔ | ✔ | ✔ | ✔ | ✔ |
| Tet F | | | ✔ | ✔ | ✔ |
| *T. striata* LAN1001 ❖ | ✔ ✪ | ✔ ✪ | ✔ ✪ | ✔ | ✔ |
| *T. suecica* ❖ | ✘ | ✘ | ✘ | ✔ | ✔ |

✔ positive PCR amplification using specific primers against algal genomic DNA
✪ Significant primer matches detected in BLASTn searches against the genome sequence.
✘ No significant primer matches detected in BLASTn searches against the genome sequence.
❖ genome  assembly available in public database

Table S4 : Sequences of qPCR primers used in this study

| **Primers** | **sequences** |
| --- | --- |
| Tsv-S2a-F | 5′-CTC ATG TAA AAC ACT TCA TTG-3′ |
| Tsv-S2a-R | 5’-CTT AAA AAA CGG ATG TGC AG-3′ |
| Tsv-S2b-F | 5′-AAT CAA CCG AGC CAG TTA CC-3′ |
| Tsv-S2b-R | 5′-TTC GAT CTG GGA ACG TAT AC-3′ |
| Tsv-S3b-F | 5′-ACA ATG TGG CGG AAG GTA AG-3′ |
| Tsv-S3b-R | 5′-CGC CGT CGA TTA GCT TTT TC-3′. |
| TetV-2-F | 5’-ATC AAC GTG GAG TTC CGG TC-3’ |
| TetV-2-R | 5’-AGGCGAGCTTGAACTTCTGG-3’ |

### Supplementary Figures :

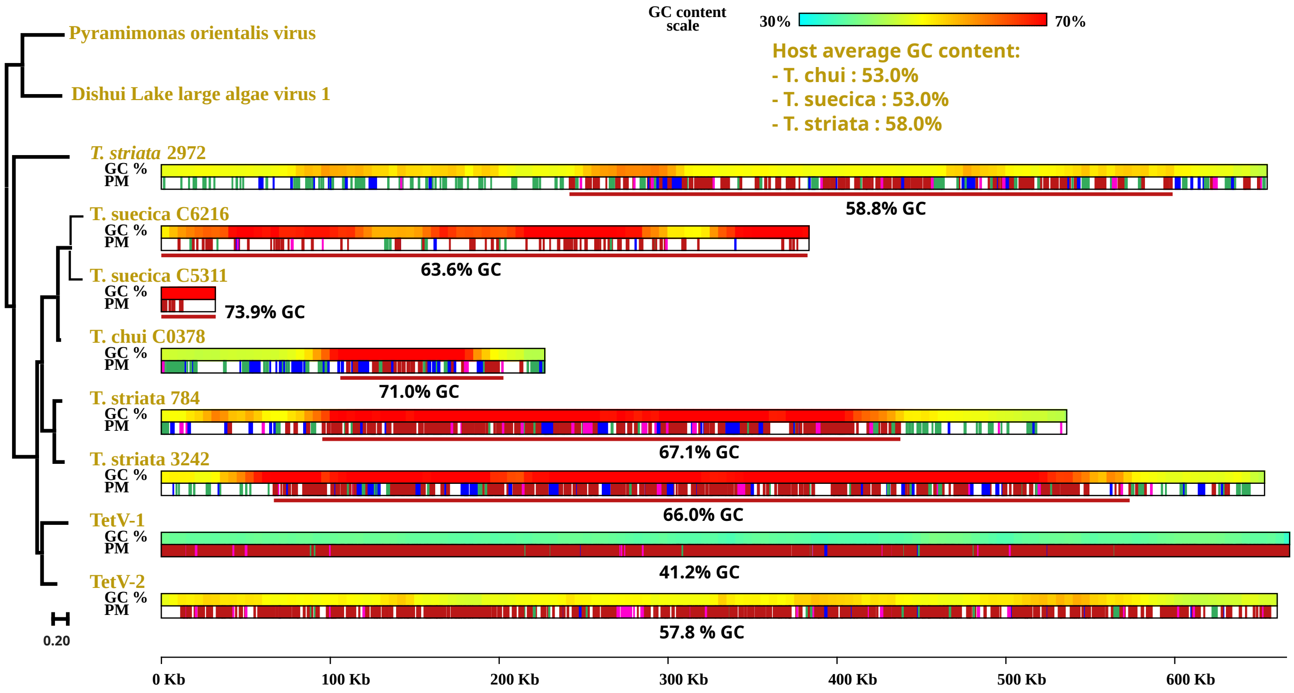


***Figure S1 | Endogenous TetV-like elements in Tetraselmis genomes.*** *Left: Phylogenetic tree of viral DNA polymerases encoded by TetV-1, TetV-2, and TetV-like insertions in Tetraselmis genomes (data from Chase et al., 2018; DOI: 10.1093/ve/veac068). The TetV-1 DNA polymerase was used as a TBLASTN query to identify homologs in algal genomes. Protein sequences were aligned with MAFFT, and phylogenetic reconstruction was performed with IQ-TREE under default parameters. All branches received >80% ultrafast bootstrap support. Right: Genomic contigs encoding viral DNA polymerases. Two color-coded tracks indicate local GC content and the taxonomic affiliation of the best BLASTX protein match in GenBank NR. Colors: red, virus; green, green alga; blue, other eukaryotes; pink, prokaryotes. Viral insertion regions are underlined in red, with the average GC content of each insertion shown below.*

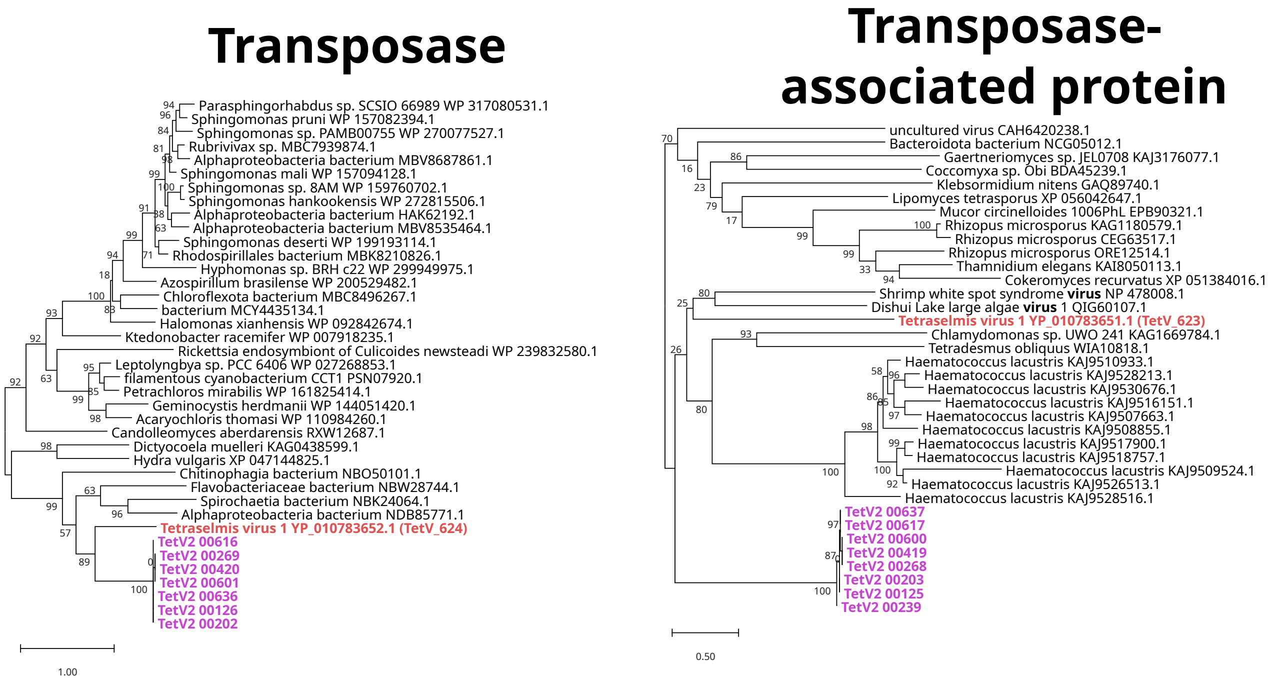


***Figure S2: phylogenetic trees of the transposases and transposase-associated proteins encoded by duplicated gene doublets.*** *Phylogenetic reconstructions were generated with FastTree using default parameters. Branch support values were estimated with the SH-like local support method implemented in FastTree.*

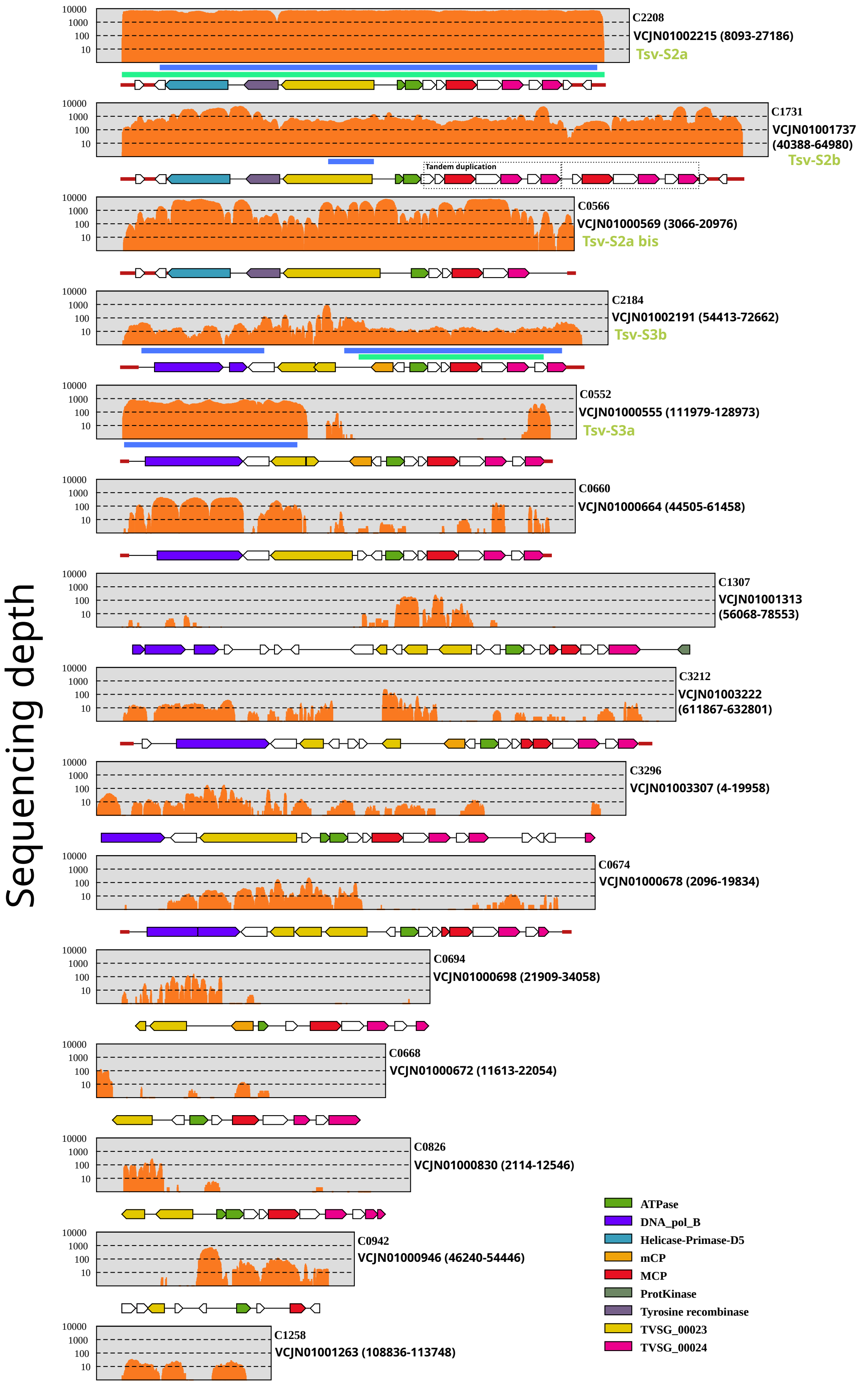


***Figure S3 : Regions of the LANL1001 genome covered by Illumina short reads..*** *Each panel shows position-specific read coverage (orange, y-axis). Below the coverage plots are the corresponding regions assembled from short reads (thick blue lines) and long reads (thick green lines), together with annotations of endogenous TetV-like elements (ETEs), including genes (colored arrows) and terminal inverted repeats (TIRs, thick red lines). Gene color codes are indicated in the figure. For each region, the region identifier, contig GenBank accession number, and genomic coordinates within the contig are provided.*

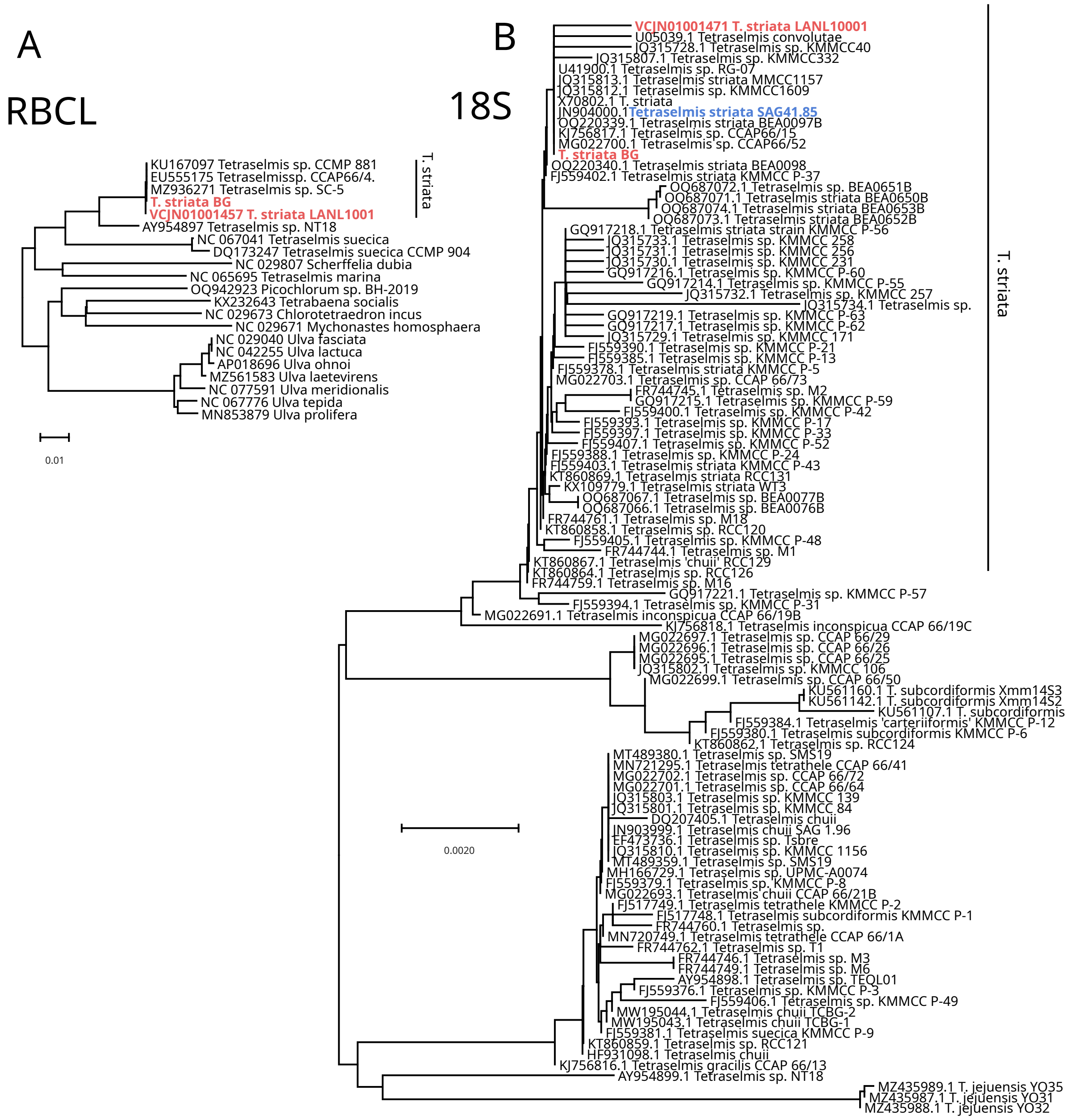


***Figure S4:*** *Minimum evolution phylogenetic trees of RBCL (A) and 18S rDNA (B) sequences generated using MEGA X.*

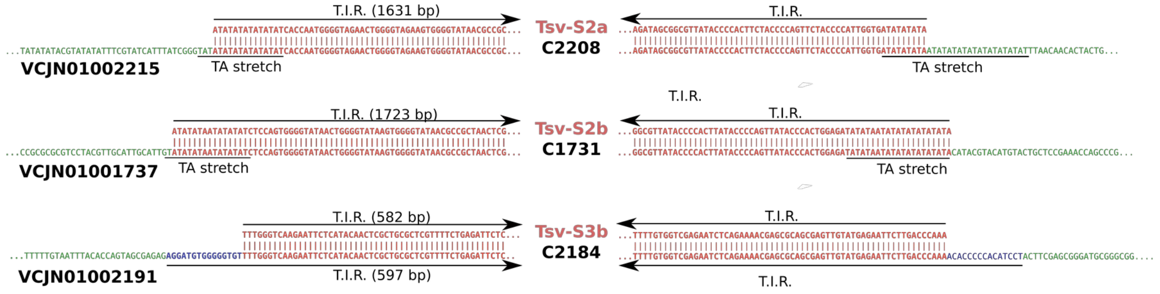


***Figure S5 : Sequence alignments at insertion sites between Tsv genomes and ETE guides.***

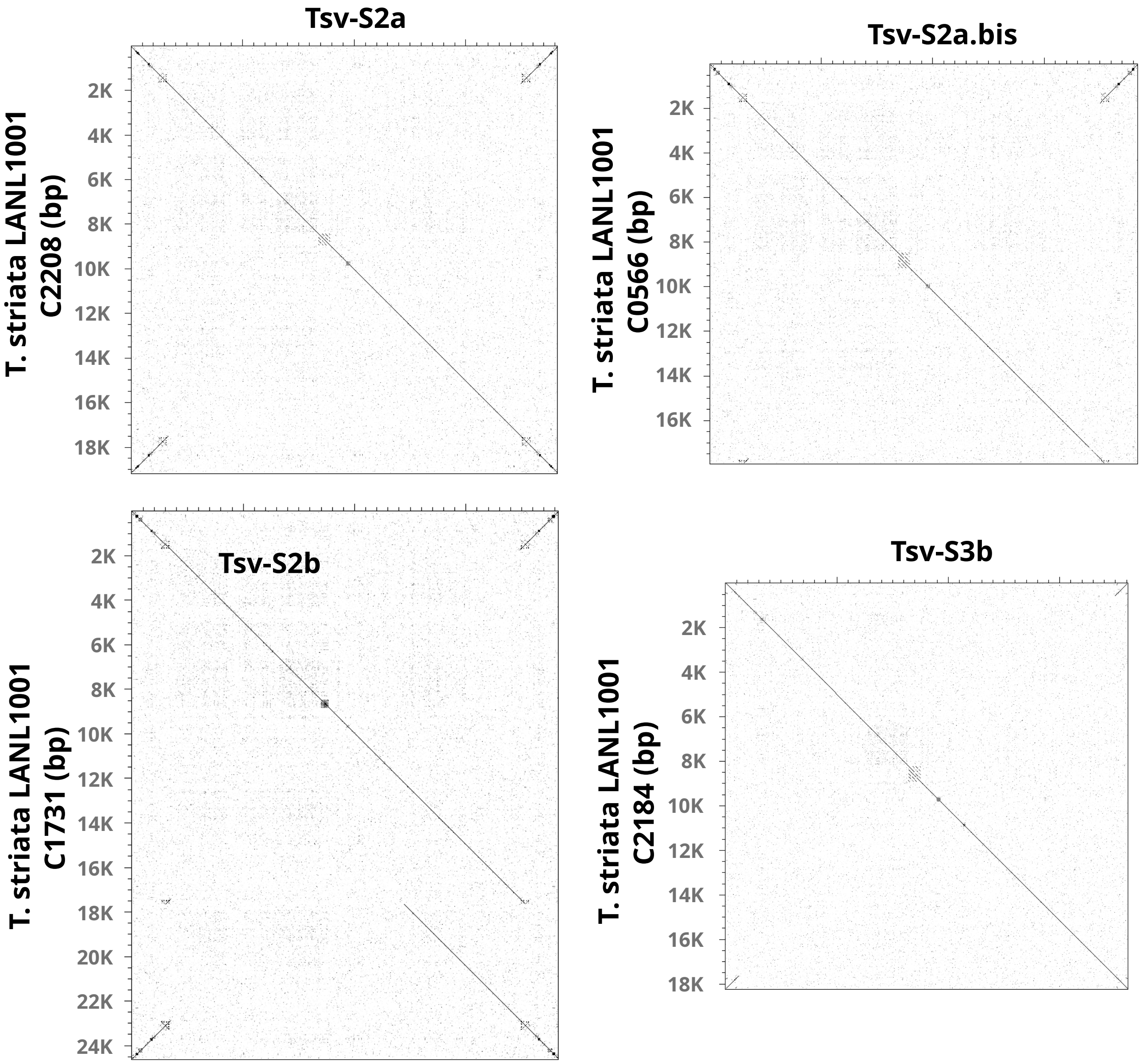


*__Figure S6__ : Alignment of assembled Tetraselmis virus (Tsv) genomes against endogenous Tsv element (ETE) reference regions used for Illumina short-read recruitment. Pairwise alignments were generated using Dotter from the SeqTools package (Sonnhammer & Durbin, 1995; Gene 167: GC1–10; doi:10.1016/0378-1119(95)00714-8).*

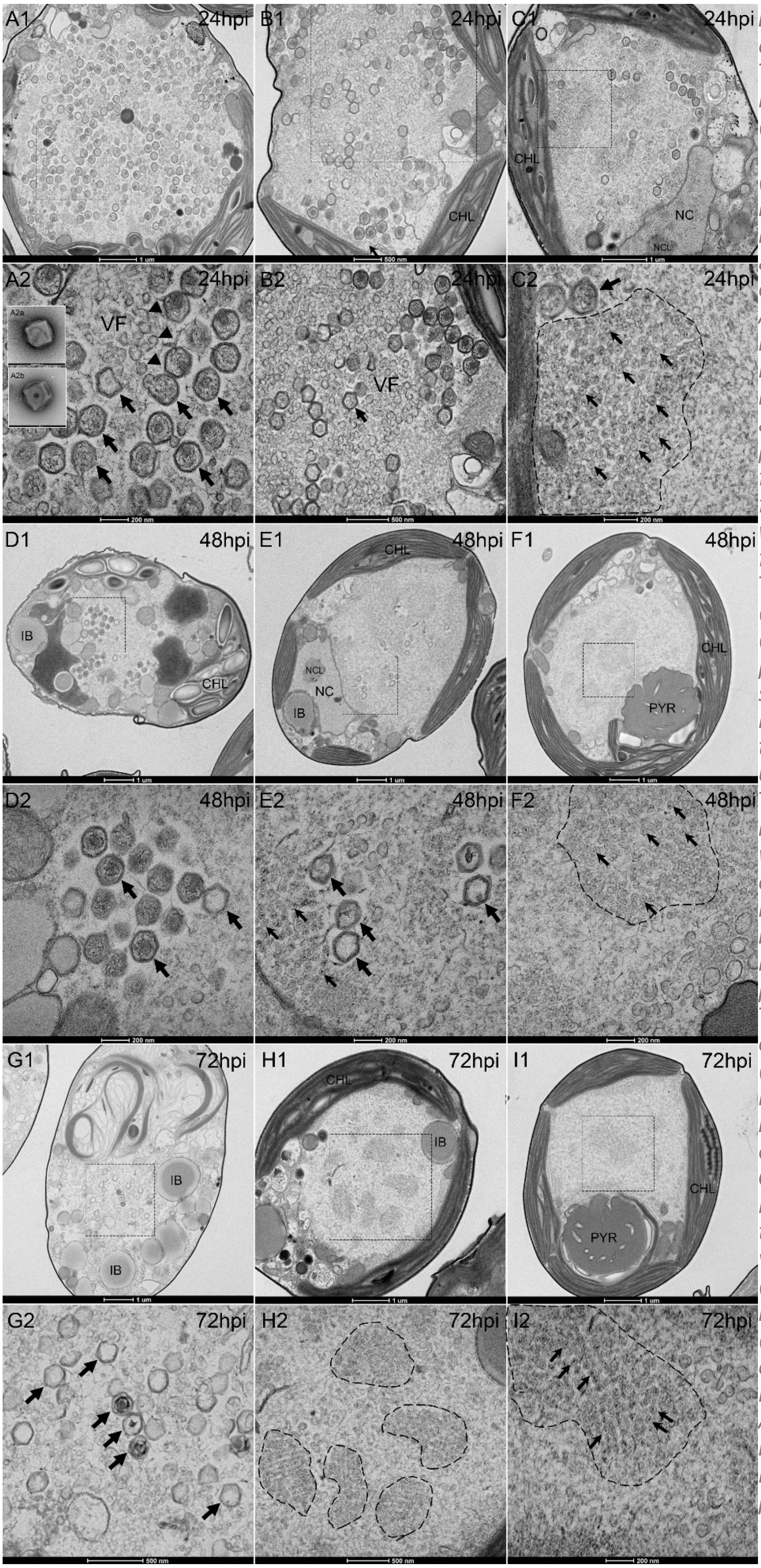


***Figure S7.*** *Transmission electron microscopy of Tetraselmis viruses during infection of Tetraselmis striata (TS).*

***(A1–C2)*** *Ultra-thin sections of infected cells at 24 hours post-infection (hpi). Panels A1, B1, and C1 show overviews of cellular organization; panels A2, B2, and C2 display higher magnifications of the boxed regions. In A2 and B2, high-magnification views reveal assembly of giant virus particles within putative viral factories (VF), with membrane fragments (arrowheads), as well as empty and partially filled particles. In C2, smaller Tetraselmis virus-like particles (Tsv) are concentrated (dashed line) at the cell periphery. Panels A2a and A2b show negatively stained mature giant virus particles at the same scale as A2.* ***(D1–F2)*** *Ultra-thin sections of infected TS cells at 48 hpi. D1, E1, and F1 show cellular overviews, while D2, E2, and F2 depict corresponding higher-magnification views. D2 and E2 reveal mature and immature empty giant virus particles, whereas F2 shows Tsv particles concentrated in a central region (dashed line).* ***(G1–I2)*** *Ultra-thin sections of infected TS cells at 72 hpi. G1, H1, and I1 show overviews of cellular ultrastructure, with G2, H2, and I2 providing higher-magnification views of the boxed areas. Most giant virus particles appear empty (G2). Peripheral and central regions rich in Tsv particles (dashed lines in H2 and I2) are evident, with Tsv assembling into pseudo-crystalline arrays. Abbreviations: NC, nucleus; NCL, nucleolus; IB, inclusion body; CHL, chloroplast; PYR, pyrenoid.*

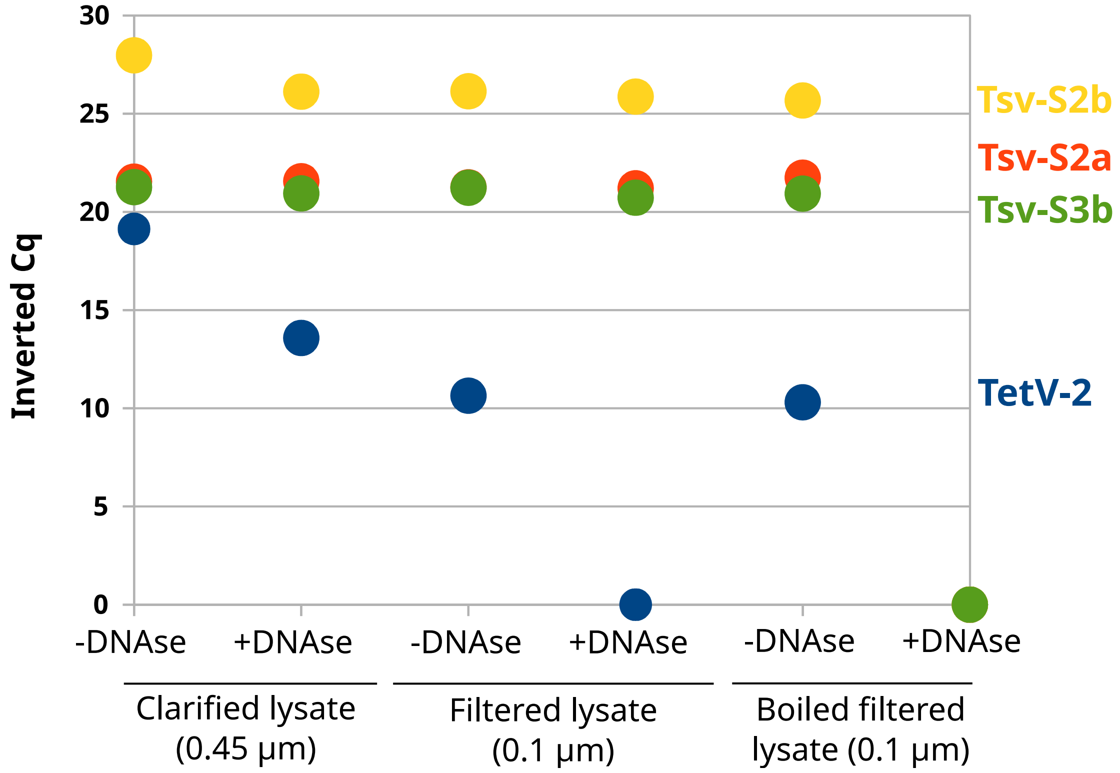


*Figure S8: qPCR assay with TetV-2 and Tsv markers.*

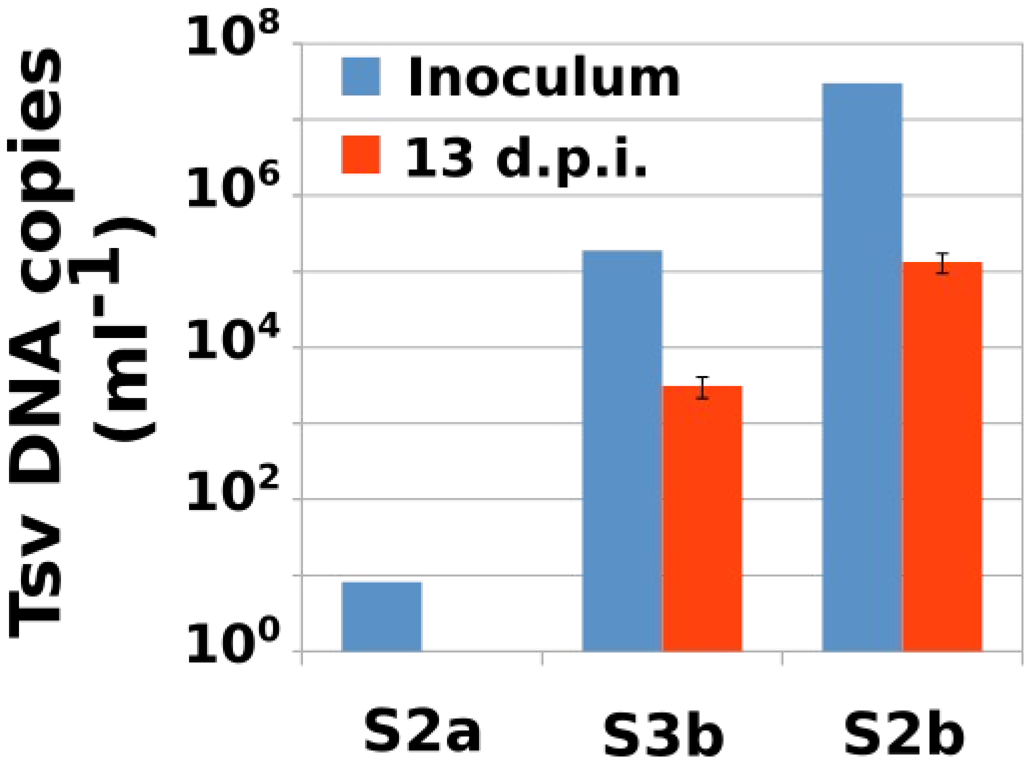


***Figure S9. Quantification of TsV DNA copy numbers at 0 and 13 days post infection (dpi).*** *TsV abundance was determined by quantitative PCR (qPCR) using strain-specific primer sets. The inoculum consisted of a TsV mixture dominated by strain TsV-S2b. Bars represent the mean values of three biological replicates, and error bars indicate the standard deviation.*

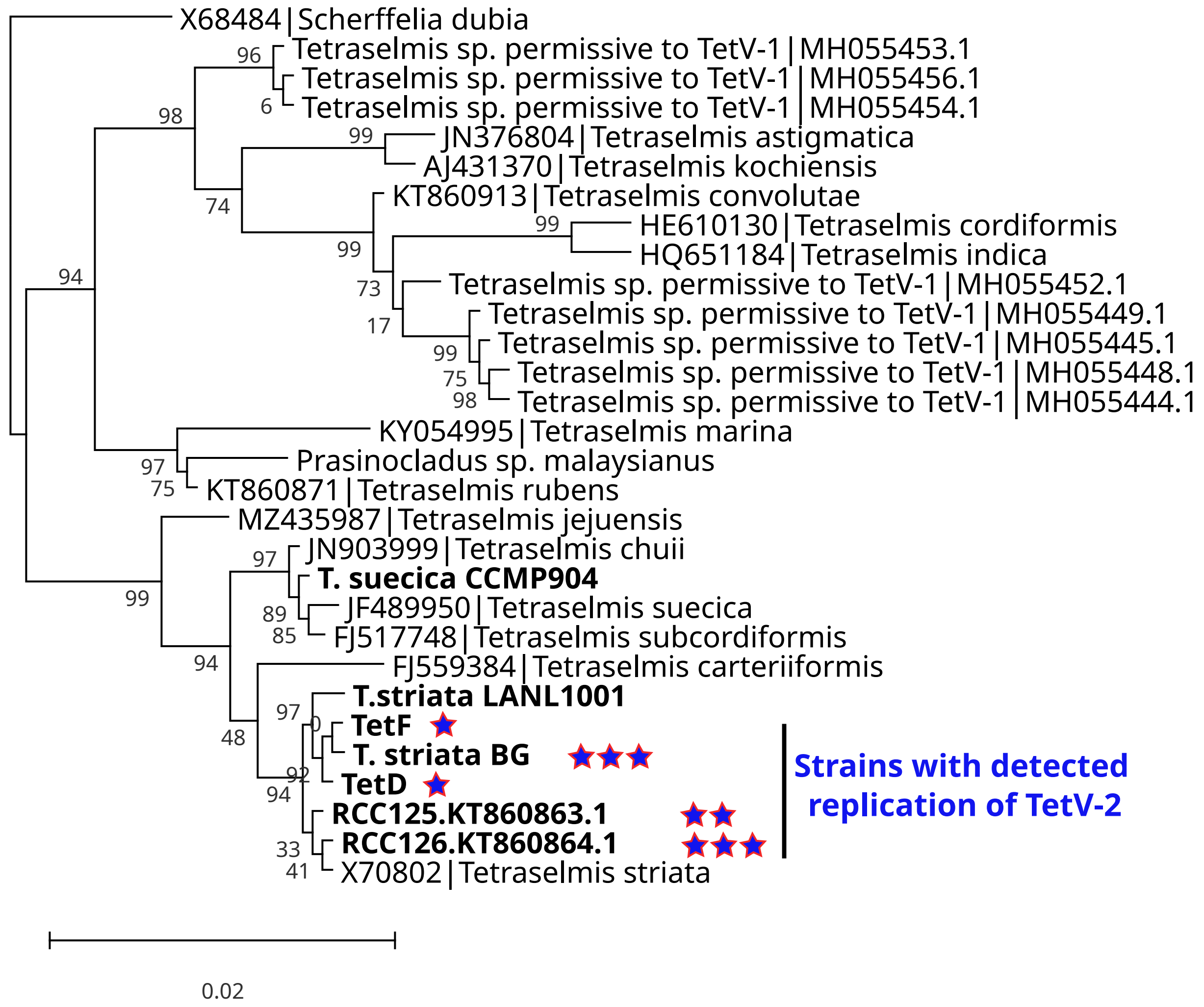


***Figure S10. 18S rRNA phylogenetic relationships among Tetraselmis strains used in the host range assay.*** *Strains tested in this study are shown in bold. TetV-2 replication levels are represented by stars, where an increasing number of stars denotes higher viral replication.*

### Supplementary Data :

Sequences of Tetraselmis striata strain GB gene markers :

>RBCL-T.striata.sp.BG

GCGTACCCTTTAGATTYATTTGAAGAAGGTTCAGTTACAAACTTATTTACTTCTATTGTAG
KTAACGTTTTTGGTTTCAAAGCTTTACGTGCTCTTCGTTTAGAAGATTTACGTATTCCAGT
TGCTTACTGTAAAACTTTTACAGGTGCTCCGCACGGTATTCAAGTTGAACGTGATAAATT
AAACAAATATGGACGTGGTTTATTAGGTTGTACTATTAAACCAAAATTAGGTCTTTCTGCT
AAAAACTACGGTCGTGCTTGTTACGAATGTTTACGTGGTGGTTTAGATTTTACAAAAGAT
GATGAAAACGTAAACTCACAAGCATTTATGCGTTGGAGAGATCGTTTCCTATTCGTATCA
GAAGCTATCTACAAATCACAAGCAGAAACTGGTGAAATTAAAGGTCACTACTTAAACGT
AACAGCAGGTACTTGTGAAGAGATGATGAAGCGTGCTGAATGTGCCGCAGGTTTCGGTG
TACCAATTGTTATGCACGATTACTTAACAGGTGGTTTTACAGCTAACACTTCATTAGCTAT
TTACTGTCGTGATAATGGTTTACTATTACACATTCACCGTGCTATGCACGCAGTTATTGAC
CGTCAACGTAATCACGGAATTCACTTCCGTGTTTTAGCAAAAGCTTTACGTATGTCTGGT
GGGGATCACCTTCACTCAGGTACTGTTGTAGGTAAATTAGAAGGTGAACGTGAAGTTAC
TTTAGGTTTCGTAGATTTAATGCGTGATGCTTACGTAGAAAAAGATCGTTCTCGTGGT

>18S-T.striata.sp.BG

aatcatgataacttcacgaatcgcatggcctccgcgccggcgatgtttcattcaaatttctgccctatcaatttgcgatggtaggatagaggcctac
catggtggtaacgggtgacggaGaattagggttcgattccggagagggagcctgagaaacggctaccacatccaaggaaggcagcaggcg
cgcaaattacccaatcctgacacagggaggtagtgacaataaataacaataccgggcttttcaagtctggtaattggaatgagtacaatctaaaca
accttaacgaggatccattggagggcaagtctggtgccagcagccgcggtaattccagctccaatagcgtatatttaagttgctgcagttaaaaa
gctcgtagttggatttcggatgggatttgccggtccgccgtttcggtgtgcactggccagtcccatcttgttgtcggggactagctcctgggcttca
ctgtccgggactaggagctgacgaggttactttgagtaaattagagtgttcaaagcaagcctacgctctgaatacattagcatggaataacatgat
aggactctggcttatcttgttggtctgtgagaccagagtaatgattaagagggacagtcgggggcattcgtatttcattgtcagaggtgaaattcttg
gatttatgaaagacgaacttctgcgaaagcatttgtcaaggatgttttcattaatcaagaacgaaagttgggggctcgaagacgattagataccgtc
ctagtctcaaccataaacgatgccgactagggattggcagacgtttttttgatgactctgccagcaccttatgagaaatcaaagtttttgggttccgg
ggggagtatggtcgcaaggctgaaacttaaaggaattgacggaagggcaccaccaggcgtggagcctgcggcttaatttgactcaacacggg
aaaacttaccaggtccagacatagtgaggattgacagattgagagctctttcttgattctatgggtggtggtgcatggccgttcttagttggtgggtt
gccttgtcaggttgattccggtaacgaacgagacctcagcctgctaaatagttactcctactttggtaggaggtgaacttcttagagggactattgg
cgtttagccaatggaagtgtgaggcaataacaggtctgtgatgcccttagatgttctgggccgcacgcgcgctacactgatgcattcaacgagcc
tagccttgaccgagaggtccgggtaatctttgaaactgcatcgtgatggggctagattattgcaattattaatcttcaacgaggaatgcctagtaag
cgtgattcatcagatcgcgttgattacgtccctgccctttgtacacaccgcccgtcgctcctaccgattgaatgtgttggtgaggagttcggattggc
agtttgtggtggttcgccactgcttacagctgagaagttctccaaaccgccccatttagaggaaggagaagtcgtaacaaggtttccgtaggtgaa
cctgc
